# Prenatal e-cigarette exposure induces region- and cell type-specific metabolic and developmental reprogramming

**DOI:** 10.64898/2026.06.19.729385

**Authors:** Wanqiu Chen, Zhong Chen, Chirag Nepal, Xue Pan, Wei-Xing Shi, Yong Li, Daliao Xiao, Wendell Jones, Malcolm Moos, Yan Dong, Charles Wang

**Affiliations:** Center for Genomics & Department of Basic Sciences, School of Medicine, Loma Linda University, Loma Linda, CA 92350; Department of Pharmaceutical Sciences, School of Pharmacy, Loma Linda University, Loma Linda, Ca 92350; Center for Perinatal Biology & Department of Basic Sciences, School of Medicine, Loma Linda University, Loma Linda, CA 92350; IQVIA Laboratories Genomics, Durham, NC 27703; Center for Biologics Evaluation and Research, Office of Cellular Therapies and Human Tissues, the U. S. Food and Drug Administration, Silver Spring, Maryland 20993; Department of Neuroscience, University of Pittsburgh, Pittsburgh, PA 15260

**Author notes:** All correspondence should be addressed to CW.

## Abstract

Prenatal e-cigarette exposure (PeCE) is increasingly prevalent and has been associated with adverse neurodevelopmental outcomes, yet how maternal vaping perturbs early brain development remains poorly understood. We integrated spatial transcriptomics, snRNA-seq and lipidomics to define neonatal rat brain responses to PeCE in rats at regional and cellular resolution. PeCE induced pronounced spatial heterogeneity in vulnerability, with the striatum exhibiting the strongest developmental transcriptional disruption. PeCE disrupts lipid metabolic homeostasis and induced molecular signatures suggestive of enhance Ca^2+^ signaling, dopaminergic responsiveness and region-specific synaptic stress programs. In the striatum, PeCE induced transcriptional programs consistent with altered lipid utilization, silent synapse-like molecular features, suppressed dendritic spine development and delayed D1-medium spiny neuron maturation. PeCE-sensitive genes were enriched in human autism spectrum disorders and neurodevelopment risk loci. Together, these findings identify disrupted metabolic and developmental reprogramming as a central feature of neonatal brain vulnerability to maternal vaping and provide a mechanistic framework linking PeCE to neurodevelopmental disease risk.

## Introduction

The use of electronic cigarettes (e-cigarettes) has increased rapidly in recent years, particularly among young adults, including pregnant women^1, 2^. Although e-cigarettes were initially promoted as a safer alternative for smoking cessation during pregnancy^3^, accumulating evidence challenges this notion. Maternal nicotine exposure is associated with significantly increased health risks in offspring, including heightened susceptibility to nicotine dependence^4, 5, 6, 7^, neurodevelopmental disorders^8, 9, 10, 11^, attention deficit hyperactivity disorder (ADHD)^12, 13, 14^, autism spectrum disorders^15^, and other psychiatric conditions^16, 17^. Despite these associations, the mechanisms by which prenatal e-cigarette exposure (PeCE) increases neurodevelopmental risk remain poorly understood.

Emerging studies indicate that PeCE induces epigenetic alterations in the developing brain, disrupts cortical neuronal development, and perturbs neurotransmitter systems and cytokine signaling pathways^18, 19^. Our recent work further demonstrates that PeCE interferes with Ca^2+^ homeostasis and alters neuronal lineage specification in the developing cortex, thereby impairing neurodevelopmental processes^20^. However, the brain is a highly heterogeneous organ composed of functionally specialized regions, and the region-specific transcriptional effects of PeCE remain largely unknown. Defining these spatially resolved molecular changes is essential for understanding the mechanisms underlying PeCE-associated neurodevelopmental disorders.

Here, we combined spatial transcriptomics, single-nucleus RNA sequencing, bulk RNA-seq, and lipidomics to define the region- and cell type-specific molecular consequences of prenatal e-cigarette exposure in the neonatal rat brain. We identified pronounced spatial heterogeneity in vulnerability across cortical and striatal regions, particularly the caudoputamen (CPu, also known as dorsal striatum in rat) emerging as a major locus of transcriptional vulnerability. PeCE induced marked dysregulation of lipid metabolic homeostasis, characterized by enhanced fatty acid and glycerolipid remodeling in the CPu and suppression of cholesterol biosynthesis programs in cortical regions. These metabolic alterations were accompanied by molecular signatures consistent with heightened Ca^2+^ influx capacity, enhanced dopaminergic responsiveness, and region-specific mitochondrial and synaptic stress adaptation. At the cellular level, PeCE promoted silent glutamatergic synapse signatures, suppressed dendritic spine developmental programs, and delayed maturation of D1 medium spiny neurons. Finally, PeCE-sensitive genes significantly overlapped human autism spectrum disorder and neurodevelopmental disorder risk loci. Together, these findings provide a spatially resolved framework for understanding how PeCE reshapes metabolic and transcriptional programming in the neonatal brain, providing a mechanistic link between maternal vaping and neurodevelopmental vulnerability.

## Results

### Study design, data generated, and quality control

We generated a spatial transcriptomic dataset of the developing rat brain following prenatal e-cigarette exposure. A chronic intermittent e-cigarette exposure rat model was used as described previously^20, 21^. Pregnant Sprague-Dawley rats were exposed to 2.4% e-cigarette vapor or control air from gestational day 4 to day 20 (**Fig. 1a**). To determine whether PeCE disrupts brain development in a region-specific manner, we analyzed spatial gene expression in coronal brain sections from postnatal day 7 (P7) offspring. Coronal sections were obtained at two anatomical planes: an anterior plane containing the nucleus accumbens (NAc), anterior cingulate cortex, and CPu, and a posterior plane containing the hippocampus, thalamus, and hypothalamus (**Fig. 1b, Suppl. Fig. 1**). In total, 34 sections (10-μm thickness) were obtained from the right hemispheres (male control, n = 6; male e-cig, n = 12; female control, n = 6; female e-cig, n = 10) and analyzed using the 10x Genomics Visium spatial transcriptomics platform (**Suppl. Fig. 2**). Across all samples, an average of 93% of reads mapped to the rat genome (rn7 assembly), with > 71% mapping to exons. Each sample yielded a mean of over 4,400 genes detected per spot (**Suppl. Data 1**), indicating high-quality spatial transcriptomic data.

**Figure 1.**
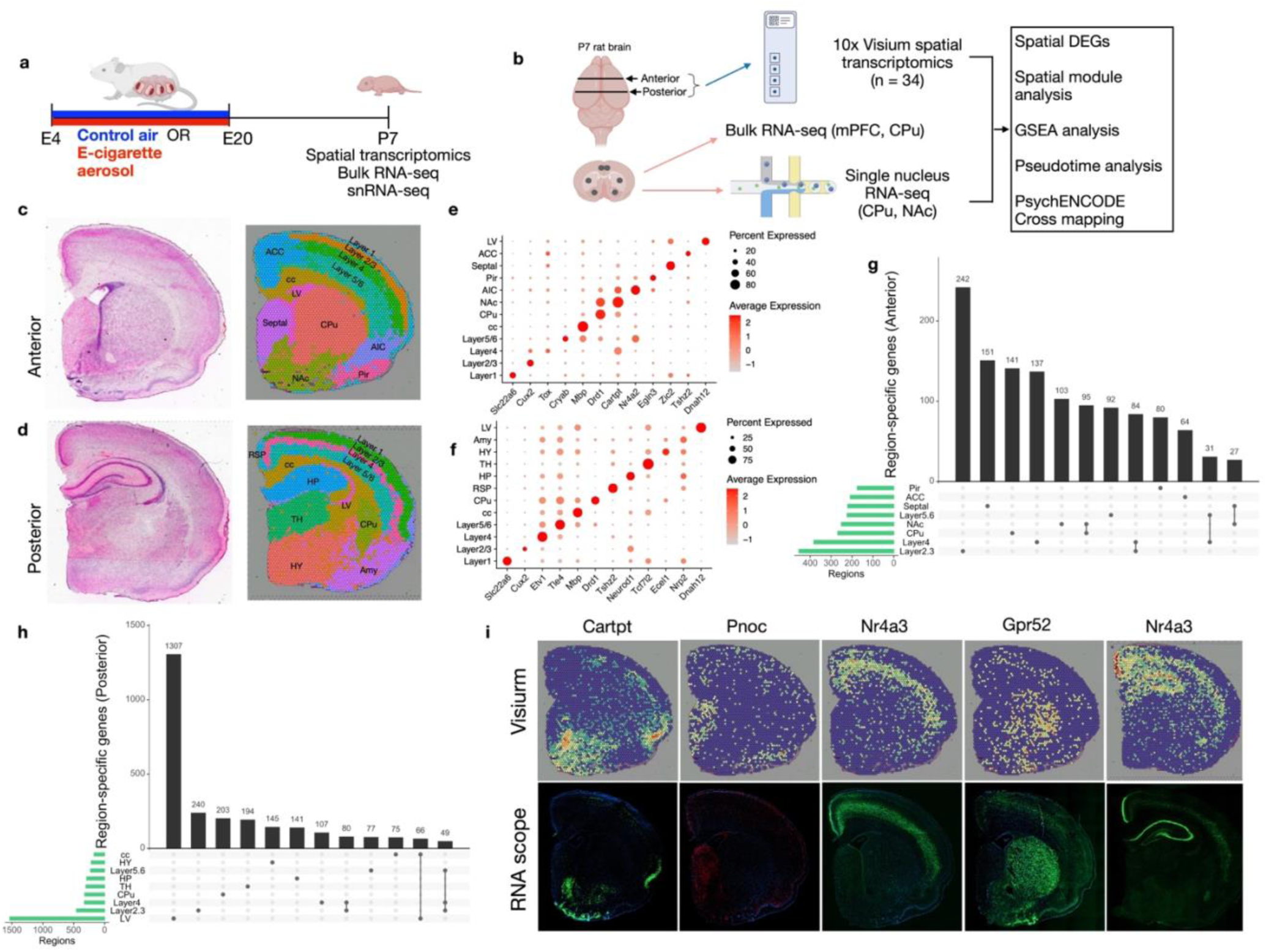
Overall study design and anatomy regions of P7 rat brain defined by spatial transcriptomics. **a,** Pregnant Sprague-Dawley rats were exposed to e-cigarette vapor from gestational E4 to E20. The brains of P7 offspring were harvested for the study. **b,** Two coronal sections (anterior and posterior) from each P7 rat brain were obtained for 10x Visium spatial transcriptomics. CPu, NAc, mPFC regions were dissected for either bulk RNA-seq or snRNA-seq. **c-d,** The left-side images are representative H&E sections of the right hemisphere anterior (c) and posterior (d). The right-side images are spatial annotations of Visium spots based on the transcriptomic signature. **e-f,** Dot plots showing the marker genes for each annotated brain region of anterior (e) and posterior (f) sections. LV, lateral ventricle; ACC, anterior cingulate cortex; Pir, piriform cortex; AIC, anterior insular cortex; NAc, nucleus accumbens; CPu, caudoputamen; cc, corpus callosum; Amy, amygdala; HY, hypothalamus; TH, thalamus; HP, hippocampus; RSP, retrosplenial cortex. **g-h,** UpSet plots showing intersections between each brain region exhibiting region-specific genes across anterior (g) and posterior (h) brain. **i,** Spatial visualization of selected region-specific gene expression in Visium spatial feature plots (top) and corresponding RNA *in situ* hybridization images by RNAScope (bottom).

The brain regions were first defined by unsupervised clustering of Visium spots based on transcriptomic signatures. The CPu and cortex exhibited the greatest number of DEGs and prominent alterations in lipid metabolism; they were therefore selected for bulk RNA-seq analysis. In parallel, the CPu and NAc were micro-dissected for single-nucleus RNA sequencing (snRNA-seq). Finally, PeCE-induced DEGs were cross-referenced with human PsychENCODE datasets^22, 23^ to assess their relevance to known neurodevelopmental disorders (**Fig. 1b**).

### Neonatal rat brain spatial transcriptomic reference map

We first integrated 17 anterior and 17 posterior spatial transcriptomic datasets, respectively, and performed unsupervised clustering analysis using the Seurat package^24^ (see Methods). This analysis identified 12 clusters in both anterior and posterior sections (**Fig. 1c-d**). Owing to a lack of a comprehensive reference atlas for the developing rat brain, we annotated anatomical regions using the Allen Mouse Brain Atlas (http://atlas.brain-map.org) as a reference, supplemented with published region-specific markers.

All 34 Visium sections showed high consistency in clustering of functional brain regions across samples (**Suppl. Fig. 3-4**) and recapitulated the anatomical organization observed in the reference atlas (Allen Mouse Brain) (**Suppl. Fig. 2a**). To define region-specific transcriptional signatures, we compared gene expression in each region against all others using a false discovery rate (FDR) <0.05 and log₂ fold-change >= 0.6, generating a region-specific enriched gene sets for each anatomical region (**Fig. 1e & 1f**, **Suppl. Data 2-3**). These gene sets exhibited minimal overlaps across regions (**Fig. 1g & 1h**), supporting their specificity as regional molecular markers. We further validated the spatial expression distribution patterns of selected markers using RNAScope *in situ* hybridization, including *Cartpt* (NAc), *Pnoc* (septal region), *Nr4a3* (cortical layer 5/6 and hippocampus), and *Gpr52* (striatum). These results were consistent with the spatial transcriptomic data, confirming region-specific gene expression patterns (**Fig. 1i**).

Region-enriched genes correspond to known neuronal populations and functional specializations. For example, *Drd1* and *Drd2* were enriched in the NAc and CPu (**Suppl. Data 2**), consistent with the predominance of dopamine D1 receptor-expressing and D2 receptor–expressing medium spiny neurons (D1-MSNs and D2-MSNs) in these regions. In addition, LIM homeobox (*Lhx*) family genes, which play pivotal roles in brain development and regional patterning, exhibited distinct spatial expression patterns. Prior studies have shown that *Lhx2* regulates the development of the forebrain^25^, *Lhx8* regulates the differentiation of cholinergic neurons^26^, and *Lhx9* regulates the thalamus development^27^. Consistent with previous reports, *Lhx2* was enriched in cortical Layer 2/3, anterior cingulate cortex (ACC), and hippocampus (HP), *Lhx8* in the NAc and hypothalamus, and *Lhx9* in the thalamus (**Suppl. Fig. 2f**).

Together, these high-resolution spatial transcriptomic profiles established a reference map of neonatal rat brain and provide a foundation for investigating how prenatal e-cigarette exposure alters brain development in a region-specific manner.

### Prenatal e-cigarette exposure induces spatially distinct transcriptomic changes in the developing brain

PeCE induces cell type-specific transcriptomic alterations^20^, but it remains unknown whether these alterations follow region-specific patterns across the developing brain. To determine the region-specific susceptibility without bias, we quantified the spatial distribution of PeCE-induced transcriptomic changes across anatomically defined brain regions using the spatial transcriptomic data shown in **Figure 1**. The MAST model^28^ (**<u>M</u>**odel-based **<u>A</u>**nalysis of **<u>S</u>**ingle-cell **<u>T</u>**ranscriptomics) was employed to overcome the challenges posed by the sparsely populated datasets commonly observed in the single cell gene expression data. At the threshold FDR of < 0.05 and log fold-change >= 0.25, we identified 2,327 unique differentially expressed genes (DEGs) in male and 2,261 DEGs in female rats in anterior sections. In posterior sections, we identified 1,116 unique DEGs in male and 1,701 DEGs in female rats **(Suppl. Data 4**). These findings suggest that PeCE exerts more pronounced effects on gene expression in anterior than posterior regions, and this is unlikely due to data variability, as the top 3,000 most variable genes exhibited comparable variance across regions (**Suppl. Fig. 5**).

To identify consistent DEGs, we focused on DEGs overlapped between male and female **(Suppl. Data 5-6**). In total, we identified 460 common DEGs in anterior, and 351in posterior. These DEGs were consistent between sexes, with comparable proportions of upregulated and downregulated genes in each brain region (**Figure 2a**). Among the anterior regions, the CPu exhibited the most DEGs (249), while in posterior regions the hypothalamus exhibited the highest DEG counts (113). These findings suggest that the CPu is the brain region most affected at P7 after PeCE. We next examined the region specificity of these DEGs by analyzing overlaps across regions with at least 40 DEGs (**Fig. 2b-c**). Approximately 50% of the DEGs were region-specific. In anterior, 145 of 249 DEGs were unique to the CPu, while 67 out of 138 DEGs were unique to the corpus callosum. In posterior, 47 of 113 DEGs were unique to the hypothalamus, and 38 of 93 DEGs were unique to the thalamus. All DEGs unique in each region are listed in **Suppl. Data 7**-**8.** These findings indicated that most DEGs induced by PeCE were region specific.

**Figure 2.**
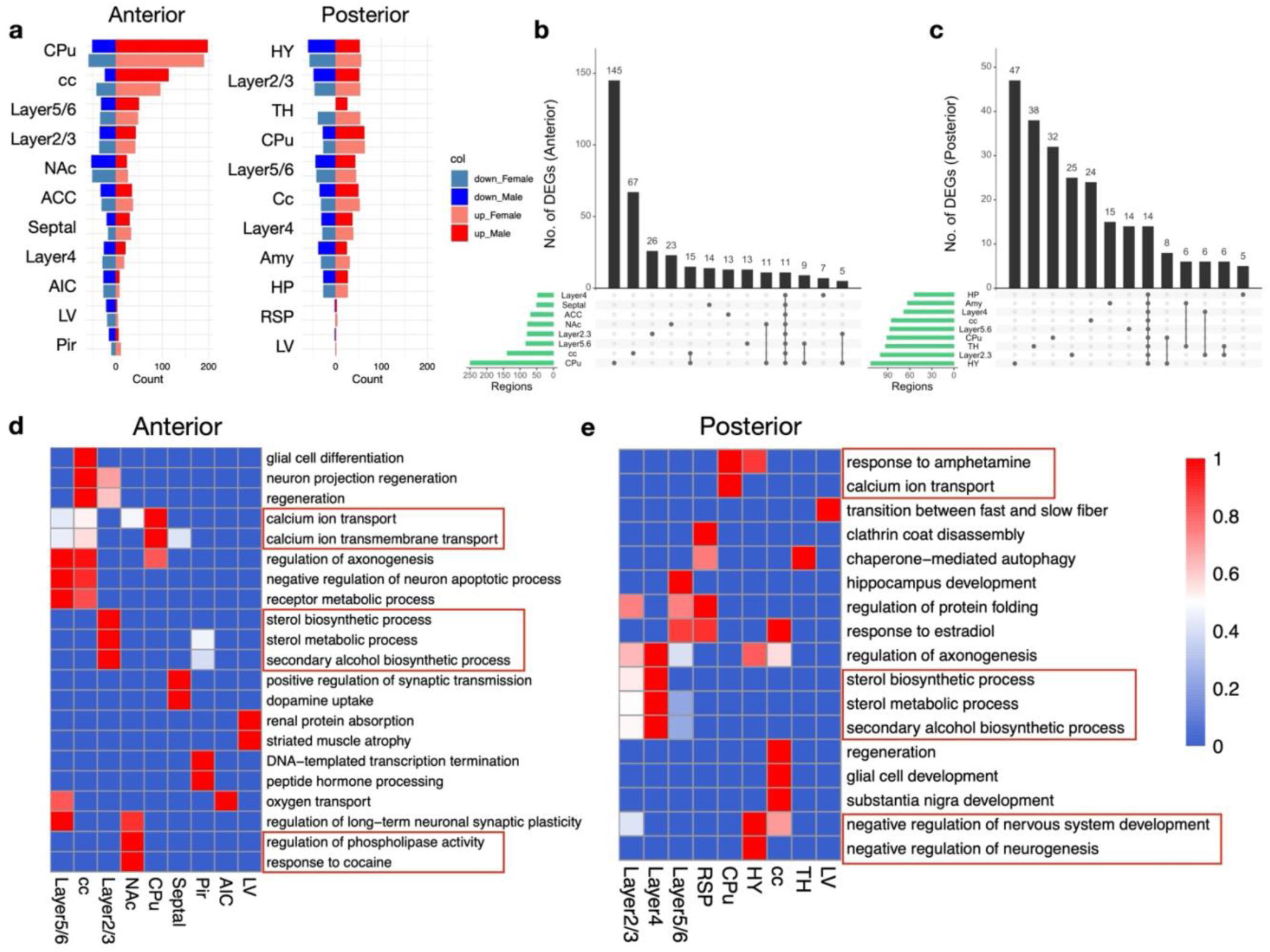
The caudoputamen transcriptome is affected most strongly at postnatal day 7 following PeCE. **a,** Number of differentially expressed genes (DEGs, Log_2_FC > 0.25 and FDR < 0.05) between e-cigarette exposed and control animals across 11 brain anterior anatomy regions (left) or 11 brain posterior anatomy regions (right) (defined by Visium datasets) at postnatal day 7, with genes up- and down-regulated shown in red and blue bars, respectively. Female, light colored bars; male, dark colored bars. **b-c,** UpSet plots showing intersections between each brain region that have more than 40 significant DEGs across anterior brain (b) and posterior brain (c). The number of DEGs from each brain region is represented by the histogram on the left (green). Dots alone indicate no overlap with any other regions. Dots with connecting lines indicate one or more overlaps of DEGs between the brain regions. The number of unique or overlapped DEGs is represented by the histogram on the top. **d-e,** Heatmap showing top enriched biological processes across individual anterior brain (d) and posterior brain (e), respectively. DEGs of each region were used for the analysis.

To elucidate the biological processes affected by PeCE, we performed Gene Ontology (GO) enrichment analysis of DEGs from each brain region (**Fig. 2d-e**). In the CPu, calcium ion and ion transmembrane transport were affected most strongly. In contrast, the corpus callosum showed enrichment in glial cell differentiation. Notably, the cortex exhibited enrichment in cholesterol biosynthesis and sterol metabolism in both anterior and posterior brain regions. These findings underscore the distinct biological pathways affected by PeCE in a brain region-specific manner.

### Prenatal e-cigarette exposure elevated molecular signatures linked to Ca^2+^ influx and dopaminergic responsiveness

Maternal smoking has been reported to increase the risk of nicotine dependence in offspring^4, 5, 6, 7^. To investigate whether PeCE alters cellular responsiveness to reward-related neurotransmission, we used spatial transcriptomics to calculate module scores, defined as the average expression of a specified gene set in individual Visium spots, for two functionally distinct pathways: the Ca^2+^ influx module and the dopamine response module. The Ca^2+^ influx module was defined using curated genes encoding the pore-forming and regulatory subunits of voltage-gated calcium channels and N-methyl-D-aspartate (NMDA) receptors, which mediate activity-dependent extracellular calcium entry. The dopamine response module comprises of key effectors of the canonical dopamine-cAMP-PKA signaling cascade. By jointly examining these calcium-entry channels and the intracellular machinery underlying postsynaptic dopamine responses, this strategy enabled high-resolution mapping of the spatial impact of PeCE on calcium and dopamine responsiveness across neonatal brain regions.

Our analysis revealed a broad increase in the Ca^2+^ influx module score following PeCE, across anterior and posterior regions (**Fig. 3a**), indicating a widespread elevation in basal Ca^2+^ influx capacity during neonatal brain development. Notably, in both male and female rats, the CPu exhibited the most robust increase in the Ca^2+^ influx module score among all regions examined (**Fig. 3b**). Consistent with this finding, the CPu also showed significant upregulation of *Atp1a1* and *Atp1a3* (**Fig. 3c**), which encode the α*-*subunits of the Na^+^/K^+^-ATPase, a critical regulator of transmembrane ionic gradients and neuronal excitability.

**Figure 3.**
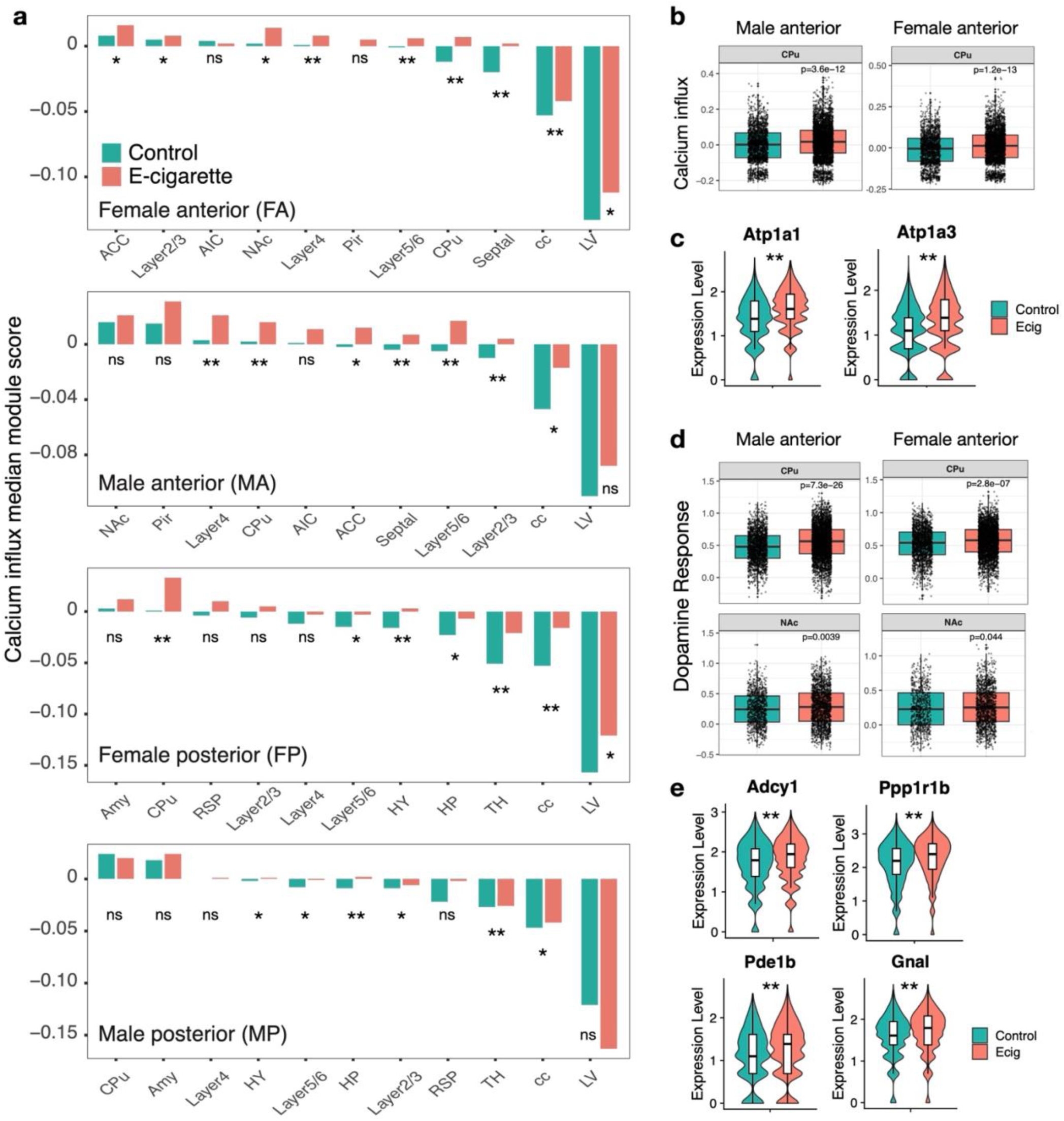
Prenatal e-cigarette exposure induces a hyper-responsive state characterized by global hyperexcitability and dopaminergic sensitization. **a,** Spatial distribution of the Ca^2+^ influx module score across anterior and posterior brain regions, demonstrating a global baseline shift in neuronal excitability. **b,** Quantitative comparison of the Ca^2+^ influx module score within the CPu. **c,** Violin plots showing significant upregulation of *Atp1a1* and *Atp1a3* in the male CPu. **d,** Boxplots showing significantly elevated dopamine response module scores in the CPu and NAc. **e,** Violin plots showing increased expression levels of key dopamine response effector genes (*Adcy1*, *Ppp1r1b*, *Pde1b*, *Gnal*) within the male CPu, highlighting sensitized dopaminergic signaling. Statistical significance for module score and gene expression was determined using the Wilcoxon rank-sum test. *p < 0.05; **p < 0.001. ns: no significant.

In addition, the dopamine response module score was likewise elevated in the CPu (**Fig. 3d**), driven by increased expression of key components of the dopamine signaling cascade, including *Adcy1*, *Ppp1r1b*, *Pde1b*, and *Gnal* (**Fig. 3e**). Together, these transcriptional changes suggest that PeCE enhances both calcium-entry capacity and dopaminergic signaling responsiveness in the CPu, potentially sensitizing the neonatal reward circuitry to subsequent environmental stimulation and leading to abnormal developmental trajectory of reward circuits.

### Prenatal e-cigarette exposure disrupts lipid metabolism in caudoputamen and cortex

On postnatal day 7, the CPu showed the highest PeCE-induced transcriptional effect, with 145 of 249 DEGs occurring exclusively in this region (**Fig. 2b**). Focusing on the CPu-specific DEGs, our GO enrichment analysis showed that genes related to lipid catabolic processes and glycerolipid/triglyceride biosynthetic pathways were most affected (**Fig. 4a**). The average expression analysis of lipid metabolism-associated genes in the CPu revealed marked upregulation following PeCE (**Fig. 4b**). To determine whether dopaminergic signaling is linked to these metabolic shifts, we evaluated the spatial co-expression patterns of *Ppp1r1b,* a key regulator of dopaminergic signaling, and the lipase *Lpl*, the rate-limiting enzyme for lipoprotein hydrolysis. We calculated Pearson correlation coefficients for this gene pair across all Visium spots and compared the strength of the associations between control and e-cigarette groups. While *Ppp1r1b* and *Lpl* exhibited a positive correlation in control rats, this correlation was intensified in PeCE rats (**Fig. 4c**). These results suggest metabolic stress and likely altered phospholipid composition in CPu after PeCE.

**Figure 4.**
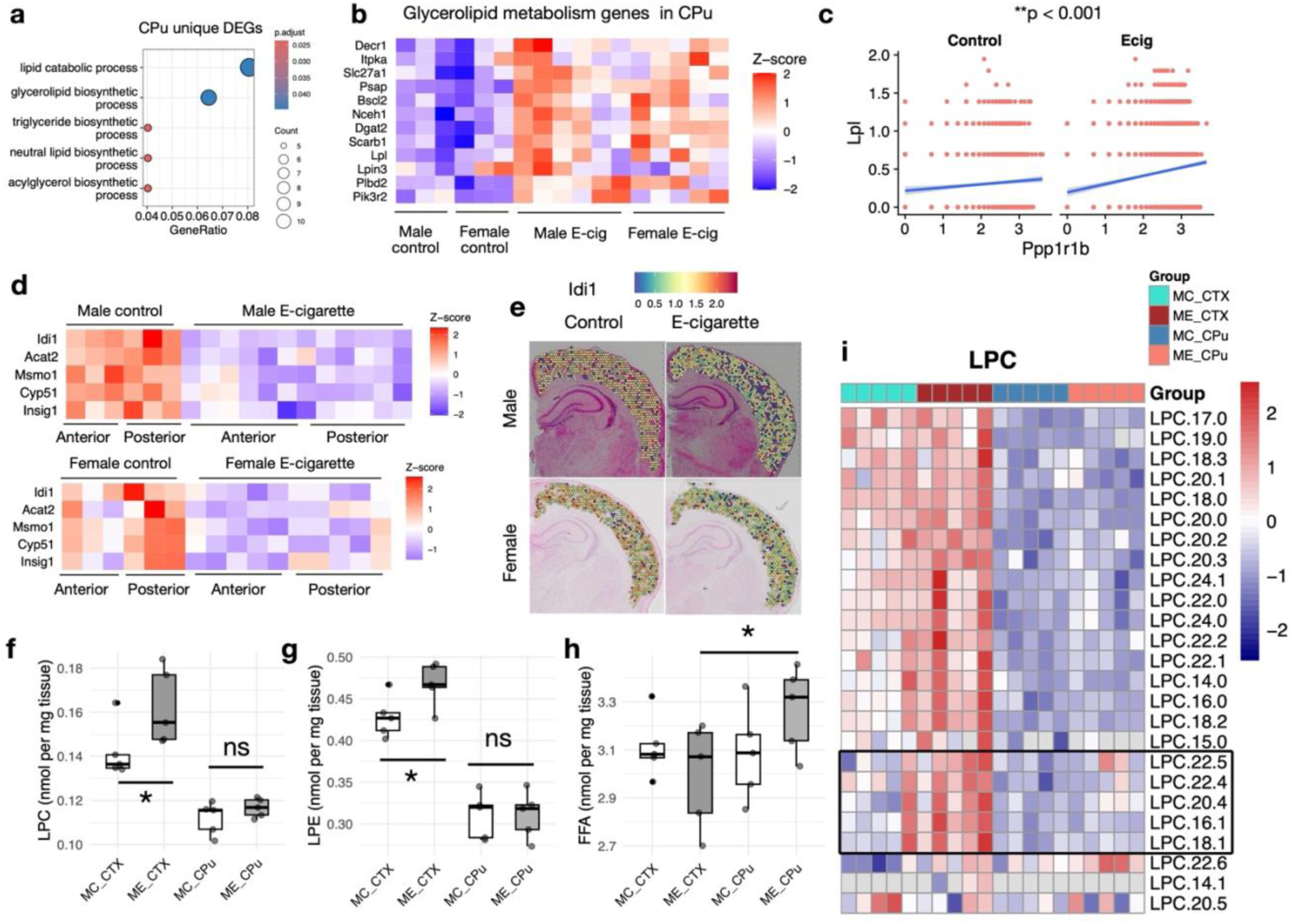
Prenatal e-cigarette exposure disrupts lipid metabolism process in the CPu and impairs cholesterol synthesis in the Cortex. **a,** Gene Ontology (GO) term enrichment analysis showing the top biological process enriched based on differentially expressed genes in caudoputamen. **b,** Heatmap showing the expression of glycerolipid metabolism-associated DEGs in CPu. **c,** Scatter plots of Lpl versus Ppp1r1b expression per Visium spot from the CPu region, split by group. Linear regression lines highlight the difference in slopes. Statistical significance of the slope difference was confirmed using a linear interaction model (** p<0.001). **d,** Heatmap showing the expression of cholesterol synthesis-related genes in the cortex region (including Layer2/3, Layer4, and Layer5/6) in both the anterior and posterior brain in each individual animal. **e,** Representative spatial feature plots showing spatial expression of *Idi1* in the posterior cortex. **f-h,** Boxplots showing total lysophosphatidylcholine (LPC) (f), lysophosphatidylethanolamine (LPE) (g), and free fatty acid (FFA) (h) quantified by shotgun lipidomics in cortex (CTX) and caudoputamen (CPu) of P7 male control (MC) and male e-cigarette (ME) rats. Each group contains five biological replicates derived from three dams. * Indicates an adjusted p-value < 0.05. **i,** Heatmap of individual LPC species measured by direct infusion mass spectrometry for P7 pups. Black box denotes changes induced by prenatal e-cigarette exposure.

In parallel, GO enrichment analysis also identified cholesterol biosynthesis and sterol metabolism as the most enriched biological processes in both the anterior and posterior cortices after PeCE (**Fig. 2d-e**), linking cortical changes to lipid metabolism as well. We therefore analyzed the average expression of cholesterol synthesis-related DEGs identified from our spatial transcriptomic dataset, including *Idi1*, *Acat2*, *Insig1, Msmo1,* and *Cyp51*, across cortical layers 2/3, 4, and 5/6 in both anterior and posterior sections. Averaged normalized counts revealed a consistent downregulation of these genes in PeCE rats (**Fig. 4d**). For instance, *Idi1*, which encodes a key enzyme in cholesterol and isoprenoid synthesis, showed marked downregulation in the posterior cortex (**Fig. 4e**). Moreover, additional region-specific cholesterol-related genes, such as *Erg28* and *Sqle* in Layer 2/3, *Fabp7* in Layer 5/6, and *Tecr* in the amygdala, were also downregulated following PeCE (**Suppl. Data 5-6**).

To validate these findings from spatial transcriptomics, we quantified approximately 970 individual lipid species in the CPu and cortex of P7 rats using a “shotgun” lipidomics approach. This analysis examined 18 lipid subclasses, including glycerophospholipids, glycerolipids, cholesterol, sphingolipids, and fatty acids^29^. Since we did not observe significant sex-dependent differences in genes related to lipid metabolism, lipidomics profiling was conducted only in male rats. In the cortex, lysophosphatidylcholine (LPC) and lysophosphatidyl-ethanolamine (LPE) were increased following PeCE (**Fig. 4f-g**), while no changes were observed in cholesterol levels. In the CPu of PeCE rats, while triacylglycerols (TGs) showed a trend toward increased levels based on saturation level and carbon chain length (**Suppl. Fig. 6**), free fatty acids were significantly elevated (**Fig. 4h**). Notably, LPC species containing unsaturated fatty acyl tails were accumulated preferentially in the cortex (**Fig. 4i**). These findings indicate that PeCE effects on lipid homeostasis in the developing brain are region-specific.

### Prenatal e-cigarette exposure induces transcriptional suppression of synaptic and dendritic programs in CPu and mPFC

Elevated levels of lysophospholipids and free fatty acids, together with stable cholesterol levels in the cortex, indicate that PeCE does not depleting bulk lipid pools but instead disrupts biosynthetic flux required for maintaining membrane-associated neuronal architectures. To determine the functional consequences of these metabolic alterations, we performed bulk RNA-seq on the CPu and mPFC followed by Gene Set Enrichment Analysis (GSEA).

In the CPu, GSEA revealed a profound metabolic stress signature characterized by strong enrichment of mitochondrial energy production pathways, accompanied by gene programs associated with reactive oxygen species (ROS) production and heightened fatty acid β-oxidation (FAO) (**Fig. 5a**, **Suppl. Fig. 7**). This transcriptional profile suggests a shift toward increased mitochondrial respiratory demand, potentially reflecting compensatory adaption to altered energy requirements. Analysis of leading-edge genes (genes contributing most significantly to the enrichment score) identified predominant enrichment of mitochondrial electron transport chain (ETC) components and FAO-related enzymes, including *Acadl*, *Acadm*, *Cpt1a*, *Ech1*, and *Hadh* (**Fig. 5b**), indicating enhanced flux of NADH/FADH2 into the ETC. Such increased electron throughput may elevate the probability of electron leakage at Complex I and III, the major sites of mitochondrial ROS generation.

**Figure 5.**
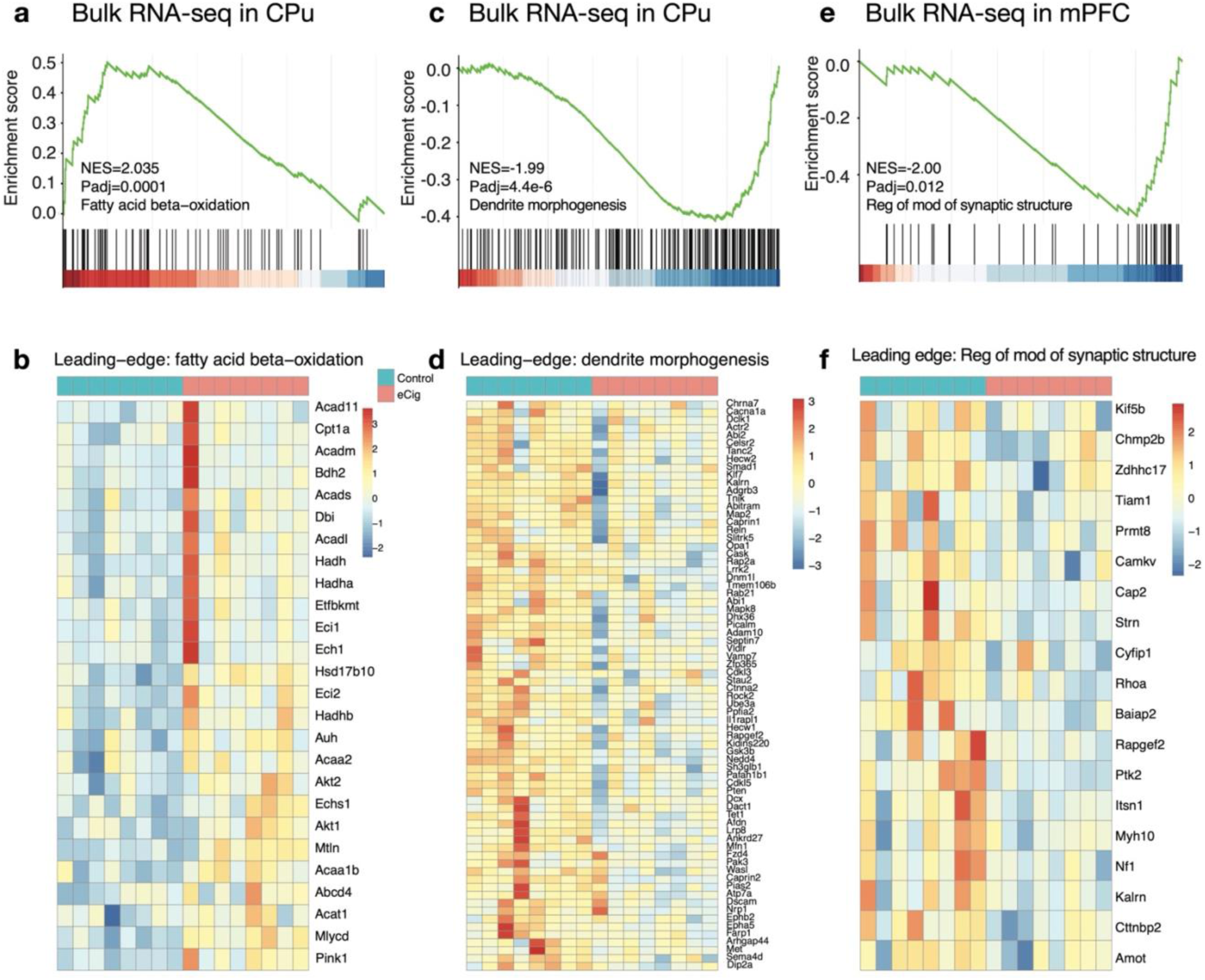
Gene set enrichment analysis (GSEA) of bulk RNA-seq data in CPu and mPFC. **a-b,** GSEA enrichment score (a) and heatmap (b) showing the PeCE-induced upregulation of genes involved in fatty acid oxidation pathway in CPu. **c-d**, GSEA enrichment score (c) and heatmap (d) showing the downregulation of genes involved in dendrite morphogenesis in CPu following PeCE. **e-f,** GSEA enrichment score (e) and heatmap (f) showing the downregulation of genes involved in synaptic structure regulation in mPFC following PeCE. NES: Normalized Enrichment Score.

Concurrently, the CPu exhibited significant downregulation of gene sets associated with neuronal structural maturation, including dendrite morphogenesis, dendritic spine development, and synaptic transmission (**Fig. 5c**, **Suppl. Fig. 7**). The leading-edge genes within these suppressed pathways included regulators of synaptic adhesion and spine morphogenesis (*Ephb2*, *Ctnna2*, *Dscam*, *Il1rapl1*) as well as actin cytoskeleton remodeling and spine dynamics (*Abi1* and *Cak15*) (**Fig. 5d)**. Together, these results indicate that PeCE induces a transcriptional state in the CPu characterized by heightened mitochondrial metabolic activity coupled with suppression of gene programs required for dendritic and synaptic development.

In contrast, the mPFC showed a more prominent disruption of synaptic structural pathways (**Suppl. Fig. 7**). Specifically, GSEA revealed significant downregulation of genes involved in synaptic organization, postsynaptic structure, actin cytoskeleton modification, and actin filament regulation (**Fig. 5e**), including leading-edge genes such as *Chmp2b*, *Strn*, *Ptk2*, *Rapgef2* (**Fig. 5f**). Although cortical cholesterol levels remained unchanged, the suppression of synaptic structural programs suggests a functional impairment of membrane-associated synaptic maintenance processes rather than a depletion of total sterol content. Given the essential role of cholesterol in synaptic membrane stability and vesicle formation, these transcriptional changes may reflect compromised synaptic maintenance and organization.

Together, these findings demonstrate that PeCE elicits a regionally distinct neurodevelopmental response across striatal and cortical regions. The CPu displays a metabolic stress-dominant state characterized by increased mitochondrial activity and concurrent suppression of dendritic developmental programs, whereas the mPFC exhibits a primary reduction in synaptic structural gene networks with comparatively limited metabolic reprogramming. These divergent transcriptional states suggest that PeCE induces region-specific adaptative responses that differentially impact energy metabolism and neuronal structural gene regulation across developing striatum and cortex.

### Prenatal e-cigarette exposure induces silent synapse molecular signatures and delays D1MSN maturation in the striatum

To determine how PeCE alters striatal neurodevelopment at cellular level, we performed snRNA-seq on CPu and NAc tissues collected at postnatal day 7 (P7). After quality control, 22,278 nuclei from the CPu and 14,414 nuclei from the NAc were retained, with an average of ∼2,300 genes detected per nucleus. Uniform manifold approximation and projection (UMAP) coupled with unsupervised clustering identified 15 transcriptionally distinct cell populations in each region (**Fig. 6a**), which were annotated using established marker genes (**Suppl. Fig. 8a**).

**Figure 6.**
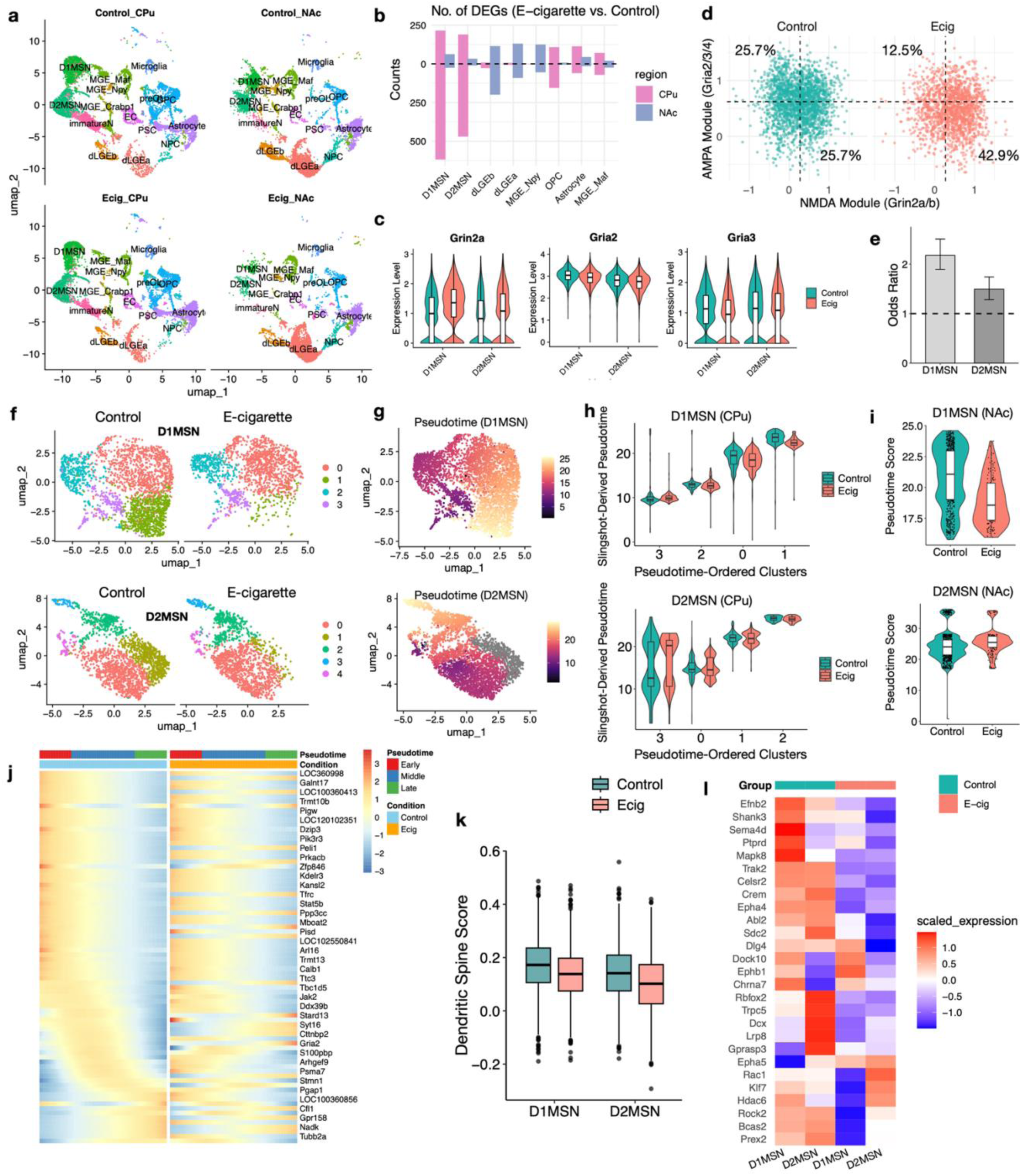
snRNA-seq analyses reveal delayed maturation of medium spiny neurons in CPu. **a,** UMAP visualization of single nuclei isolated from CPu and NAc of P7 rats following prenatal e-cigarette or control air exposure. **b,** Number of differentially expressed genes (DEGs, Log_2_FC > 0.3 and p.adj < 0.05) between e-cigarette exposed and control animals across all cell types, with up-regulated genes shown above y = 0 and down-regulated genes shown below y = 0. **c,** Violin plots showing the expression level of *Grin2a, Gria2* and *Gria3* in the CPu D1MSN and D2MSN. **d,** Scatter plot showing the module score of AMPA and NMDA of CPu D1MSN in four quadrants. **e,** Analysis of NMDA^high^/AMPA^low^ cell proportion in e-cigarette CPu MSN compared with control. The odds ratio was computed using Fisher’s exact test. **f,** UMAP visualization of single nuclei from D1MSN (top) and D2MSN (bottom) subclusters after re-clustering. **g.** Pseudotime progression derived from Slingshot analysis, illustrating the inferred developmental trajectory of D1MSN (top) and D2MSN (bottom). **h.** Violin plots showing the distribution of Slingshot-derived pseudotime value for subcluster 0-3 in UMAP (e). *p < 0.001 (Wilcoxon rank sum test). **i.** Violin plots showing the distribution of Slingshot-derived pseudotime value for NAc D1MSN (top) and D2MSN (bottom). *p < 0.001 (Wilcoxon rank sum test). **j.** Heatmaps in control and e-cigarette panels showing the expression patterns of p-value top-ranking significant genes (rows) by pseudotime-ordered CPu D1MSN (columns). tradeSeq fitGAM model: conditionTest method. **k.** Boxplot showing the module score of the CPu D1MSN and D2MSN clusters with respect to the dendritic spine formation signature. *p < 0.05, Wilcoxon rank-sum test. **l.** Heatmap showing the average gene expression of the CPu D1MSN and D2MSN enriched in the dendritic spine formation pathway.

Differential expression analysis revealed that the most prominent PeCE-induced transcriptional changes occurred in D1 and D2 MSNs within the CPu **(Fig. 6b**). In D1MSNs, 215 genes were upregulated and 618 were downregulated (Log_2_FC > 0.3; Bonferroni-adjusted p-value < 0.05). Among these DEGs, the NMDA receptor subunit *Grin2a* was increased in both D1MSN and D2MSN, while AMPA receptor subunits *Gria2* and *Gria3* were reduced (**Fig. 6c**). This reciprocal shift in glutamate receptor composition suggests transcriptional bias toward NMDA^high^/AMPA^low^ excitatory synapses, a molecular configuration characteristic of silent synapse states during early brain development^30^.

To quantify this synaptic immaturity signature, we calculated the AMPA receptor module score (*Gria2/3/4*) and NMDA receptor module score (Grin2a/b) for individual CPu D1MSNs and classified all the D1MSNs into four quadrants according to relative module enrichment (**Fig. 6d**). PeCE resulted in a 17% increase in the proportion of D1MSNs exhibiting an NMDA^high^/AMPA^low^ profile, consistent with key features of glutamatergic silent synapses (**Fig. 6d**). Similar analysis of CPu D2MSNs showed a more modest enrichment of this profile (**Suppl. Fig. 8b**). Accordingly, the odds ratios for increased NMDA^high^/AMPA^low^ nuclei following PeCE were 2.17 in D1MSMs and 1.49 in D2MSNs (**Fig. 6e**), indicating preferential vulnerability of D1MSNs to glutamatergic immaturity.

Consistent with impaired neuronal developmental progression, KEGG enrichment analysis of downregulated genes identified axon guidance as a significantly suppressed pathway in both D1MSNs and D2MSNs (**Suppl. Fig. 8c-d**). Independent re-clustering of D1MSN and D2MSN populations identified four D1MSN subclusters and five D2MSN subclusters in both control and PeCE animals (**Fig. 6f**). For D1MSNs, pseudotime trajectory analysis positioned subclusters 0 and 1 at more advanced maturation states (**Fig. 6g**). Notably, both mature D1MSN subclusters were selectively reduced in the CPu following PeCE (**Fig. 6h**), indicating delayed D1MSN developmental progression. A similar reduction in Slingshot-derived pseudotime progression was also observed in D1MSNs from the NAc (**Fig. 6i**). In contrast, D2MSNs from either CPu or NAc exhibited minimal changes in pseudotime trajectory between control and PeCE groups (**Fig. 6h-i**), suggesting that the maturational delay is predominantly restricted to the D1 lineage.

To further define the transcriptional dynamics underlying this developmental impairment, we applied tradeSeq^31^ to model gene expression changes along CPu D1MSN pseudotime using negative binomial generalized additive models. This allowed us to identify genes with significantly altered temporal expression dynamics during D1MSN maturation in PeCE rats compared to controls. PeCE disrupted D1MSN maturation in the CPu, as genes that increased during early or intermediate pseudotime in control rats exhibited delayed upregulation, shifting toward late pseudotime in PeCE rats (**Fig. 6j**). Delayed genes included multiple regulators of synaptic organization and neuronal differentiation, such as *Cttnbp2*, *Gria2*, *Psma7*, *Stmn1*, and *Arhgef9,* further supporting a retardation of the normal D1MSN maturation program.

Integrating of these single-cell findings with the bulk RNA-seq suggested reduced dendrite morphogenesis and MSN development, led us to test whether PeCE suppresses transcriptional programs required for MSN spine formation and stabilization. We therefore curated a 27-gene dendritic spine development module and computed module scores for individual MSN nuclei. Both D1MSNs and D2MSNs exhibited significantly reduced dendritic spine module scores following PeCE (**Fig. 6k**). Several genes showed pronounced reductions in D1MSNs, including *Efnb2*, *Shank3* and *Ptprd*, which encode postsynaptic scaffolding and adhesion molecules required for stabilizing nascent spines and promoting the transition from filopodia-like to mushroom-like spines. Additional reductions were observed in genes regulating actin remodeling through Rho-family GTPase signaling (*Epha4*, *Rock2*) and in genes supporting the metabolic and transcriptional machinery of dendritic spine maturation (*Trak2*, *Crem*) in both D1MSNs and D2MSNs (**Fig. 6l**).

Together, these findings demonstrate that PeCE induces a coordinated striatal immature transcriptional state characterized by silent synapse-like features, suppression of dendritic spine developmental program, and delayed D1MSN maturation in early postnatal neonatal brain.

### Prenatal e-cigarette exposure-induced transcriptional perturbations overlap human neurodevelopmental disorder risk genes

To investigate the association between PeCE-induced DEGs and human neurodevelopmental diseases, we performed the Fisher’s exact test to assess the statistical significance of overlap between P7 spatial transcriptomics DEGs and neurodevelopmental disorder-associated genes in the PsychENCODE dataset^32, 33^ and other human autism spectrum disorder (ASD) and neurodevelopmental delay (NDD) associated gene sets^34, 35^ (**Fig. 7a**). Our analysis revealed a significant overlap between P7 DEGs in rats and genes implicated in human ASD (**Fig. 7b**, **Suppl. Data 9**), suggesting that PeCE engages molecular pathways that may confer susceptibility to neurodevelopmental disorders. We next assessed the regional specificity of this association by quantifying overlaps between orthologous PeCE-induced DEGs and ASD/NDD risk genes across brain areas. The CPu exhibited the highest overlaps (**Fig. 7c**), indicating that PeCE elicits transcriptional perturbations in striatal circuits that are similar in human ASD and NDD.

**Figure 7.**
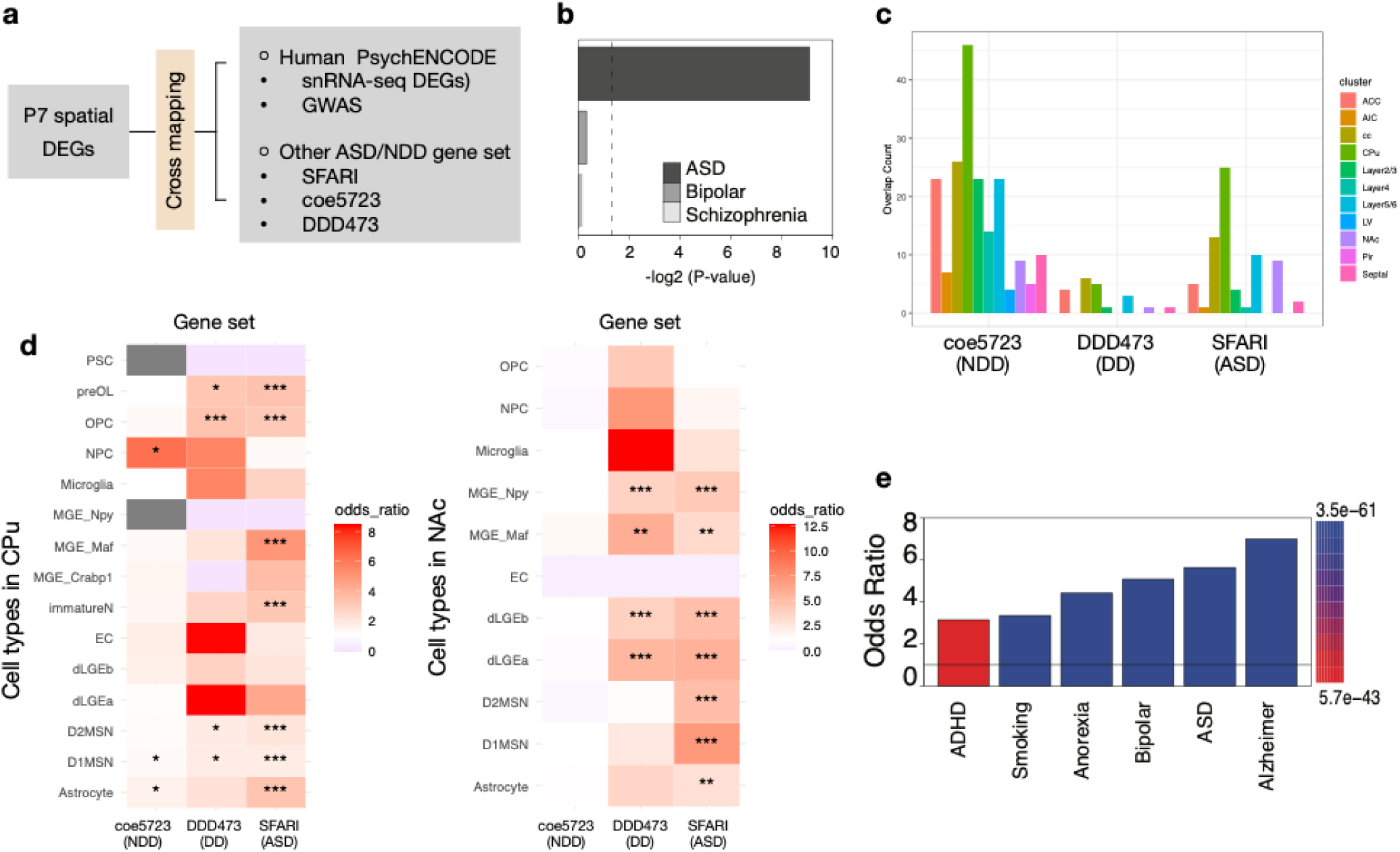
Cross mapping of prenatal e-cig exposure-induced differentially expressed genes with gene sets implicated in human neurodevelopmental diseases. a, Schematic illustration of the workflow for mapping prenatal e-cig induced DEGs from postnatal day 7 (P7) to human PsychENCODE and other ASD/NDD gene sets. b, Enrichment analysis showing overrepresentation of e-cig induced DEGs versus non-DEGs in rat compared with the orthologous genes associated with autism spectrum disorder (ASD), bipolar disorder, and schizophrenia gene sets. The x-axis shows -log2(P-value) with greater values indicating stronger statistical significance. P-values were computed using Fisher’s exact test. c, Spatial distribution of DEG number of e-cig induced DEGs overlapped with genes associated with ASD and NDD (SFARI: https://gene.sfari.org/database/human-gene/, DDD473^35^, coe5723^34^). d. Enrichment of DEGs from snRNA-seq in CPu (left) and NAc (right) that overlap with human orthologous genes associated with ASD and NDD, including genes enriched in de novo non-coding mutations. P-values were computed using two-sided Fisher’s Exact test. Asterisks, Bonferroni-corrected significance. ***p < 0.001, **p<0.01, *p<0.05. e, Enrichment analysis of human psychiatric GWAS loci orthogonally mapped to rat DEGs compared with non-DEGs were significantly enriched on DEGs. The odds ratio was computed using Fisher’s exact test.

To resolve these associations at cellular resolution, we next examined whether cell-type-specific DEGs were enriched in human ASD- and NDD-associated orthologous genes, including genes harboring *de novo* non-coding mutations. DEGs from both the CPu and NAc had significant enrichment across most major cell types within the SFARI ASD gene set (**Fig. 7d**). Consistent with these findings, multiple GWAS single nucleotide polymorphisms (SNPs) compiled by PsychENCODE were enriched in the DEGs compared with non-DEGs in P7 rats (**Fig. 7e**), where datasets related to Alzheimer’s and ASD showed the highest enrichment. Collectively, these results demonstrate that PeCE-induced DEGs, particularly within striatal regions, are enriched for genetic signatures of human neurodevelopmental disorders, underscoring the biological relevance and potential translational impact of prenatal e-cigarette exposure.

## Discussion

We generated the first spatial transcriptomic atlas of the developing rat brain following PeCE and identified pronounced spatial heterogeneity in molecular vulnerability across neonatal brain regions. Among these, the caudoputamen emerged as a major locus of transcriptional vulnerability. PeCE induced broad molecular signatures suggestive of enhanced Ca^2+^ signaling and dopaminergic responsiveness, together with region-specific perturbations in lipid metabolic homeostasis. Integrative analyses further revealed suppression of synaptic and dendritic developmental gene programs, delayed maturation of striatal medium spiny neurons (D1MSNs), and significant overlap between PeCE-sensitive genes and human neurodevelopmental disorder risk loci. Collectively, these findings provide a spatially resolved molecular framework through which maternal vaping may disturb early brain circuit assembly.

One of the most prominent responses to PeCE was the coordinated upregulation of genes involved in Ca^2+^ and dopaminergic signaling in the neonatal brain. Previous studies have shown that nicotine exposure and withdrawal increase neuronal excitability across multiple brain regions, including hippocampal CA1 pyramidal neurons, interpeduncular nucleus GABAergic neurons, and cortical neurons^36, 37, 38^, as well as Ca^2+^ activity in astrocytes^39^. In our study, we observed global upregulation of voltage-gated calcium channel and NMDA receptor genes, and upregulation of dopaminergic signaling regulators in CPu and NAc. This calcium-associated transcriptional signature was accompanied by coordinated upregulation of Ca^2+^/calmodulin-activated adenylyl cyclase *Adcy1*, and key dopamine-cAMP signaling regulators, including*Ppp1r1b*, *Pde1b*, *Gnal*, and *Rgs9*. These coordinated changes suggest an enhanced dynamic responsiveness of dopamine-cAMP signaling pathways to excitatory and dopaminergic inputs. These PeCE-induced signaling perturbations strongly converge on pathways implicated in neurodevelopmental disorders. Dysregulation of Ca^2+^ signaling is a hallmark of neurodevelopmental disorders^40, 41^. Several Ca^2+^-related genes altered by PeCE, including *Cacna1a*, *Grin2b*, *Grin2a*, *Camk2b*, *Shank3*, overlap established risk loci for ASD, epilepsy, intellectual disability, and ADHD^42, 43, 44, 45^. In parallel, substance use disorders are associated with dysregulation of cAMP signaling^46^. The upregulation of *Adcy1* and *Ppp1r1b* observed here mirrors molecular adaptions reported following chronic drug exposure^46^. Notably, ADCY1 overexpression has been linked to aberrant synaptic plasticity ^47^, abnormal cortical excitability ^48^, enhanced autism-related behavior and reduced sociability^48, 49^. Together, these observations suggest that PeCE engages transcriptional networks that intersect both activity-dependent neurodevelopmental pathways and reward-related signaling programs, raising the possibility that prenatal vaping sensitizes neonatal brain circuitry along axes relevant to later neuropsychiatric vulnerability.

A key finding of this study is that PeCE induces marked region-specific disruption of lipid metabolic homeostasis. Previous studies have shown that smoking disrupts lipid metabolism^50, 51^. However, how PeCE reshapes the lipid homeostasis in the developing brain remains unknown. In the CPu, genes associated with glycerolipid metabolism and lipoprotein hydrolysis were upregulated, accompanied by enhanced fatty acid β-oxidation signatures, increased mitochondrial respiratory pathway enrichment, and elevated free fatty acid levels. These coordinated changes are consistent with a shift toward lipid-derived energy utilization and increased oxidative metabolic demand. Such metabolic remodeling may increase developmental vulnerability of striatal circuitry to later perturbations associated with addiction-related behaviors^52,53^. In contrast, cortical regions exhibited suppression of mevalonate/cholesterol biosynthesis genes, including *Idi1*, *Msmo1*, *Sqle*, *Cyp51*, and *Dhcr7* and *Insig1*, and *Acat2*, consistent with prior studies of chronic nicotine exposure^54, 55^. Lipidomics revealed elevated LPC and LPE, indicating increased membrane phospholipid degradation and turnover. Although cholesterol levels remained unchanged, GSEA from the mPFC suggested impaired synaptic and structural remodeling. Together, these findings suggest that the cortex undergoes heightened membrane phospholipid turnover yet fails to engage in biosynthetic scaling required for active circuit assembly during early development. Given the essential role of cholesterol in synaptogenesis, myelination, and neuronal signaling^56^, and the fact that neonatal neurons rely heavily on autonomous cholesterol synthesis for axon outgrowth and synapse formation^56^, disruption in cholesterol-related pathways following PeCE is likely to impair cortical neuronal maturation and circuit development. Rather than representing a uniform metabolic injury, PeCE appears to induce distinct adaptive states in striatal and cortical regions, reflecting differential energetic and biosynthetic pressures across developing neuronal systems. Such perturbations may increase susceptibility to neurodevelopmental and psychiatric disorders, including ASD^57, 58, 59, 60^, schizophrenia^61, 62, 63^, and ADHD^64, 65^,

Single-nucleus transcriptomic analysis further points this developmental vulnerability to medium spiny neurons, particularly D1MSNs. PeCE increased the proportion of D1MSNs and D2MSNs exhibited an NMDA^high^/AMPA^low^ ^66^ transcriptional signature consistent with silent synapse-like immature glutamatergic states^67^, while pseudotime analysis revealed delayed progression of D1MSNs along their normal maturation trajectory. In parallel, genes involved in dendritic spine formation, including *Efnb2* and *Shank3,* which are crucial for establishing and maintaining stable glutamatergic synapses, were reduced by PeCE. We postulate that this aberrant synapse and MSN development is linked to the metabolic shift observed in the CPu. Developing MSNs may prioritize lipid-derived energy production to support nicotine-induced hyperexcitability at the expense of biosynthetic processes required for synapse formation. In other words, developing neurons exposed to PeCE undergo a metabolic allocation shift in which resources are increasingly directed toward sustaining heightened energetic demand at the expense of biosynthetic processes required for structural maturation. Although direct metabolic flux measurements were not performed, these observations support a model in which altered lipid utilization may compete with biosynthetic processes required for dendritic spine maturation and stable synaptic assembly. Diverting lipid resources toward beta-oxidation and ATP synthesis may compromise the availability of stable lipid-raft scaffolds^68^ necessary for synaptic maturation and dendritic spine formation. Thus, our data support a model in which PeCE creates an imbalance between metabolic adaptation and neuronal growth, resulting in region-specific suppression of developmental programs necessary for normal circuit refinement. Because silent synapses and impaired MSN maturation have been implicated in neurodevelopmental disorders like autism-related circuit dysfunction and intellectual disability^69, 70^, the striatum may represent a particularly sensitive developmental target of prenatal e-cigarette exposure.

Consistent with this interpretation, cross-species integration revealed significant enrichment of PeCE-induced DEGs within human ASD, neurodevelopmental delay, and psychiatric disease-associated gene sets, with the strongest overlap observed in CPu-responsive genes. Furthermore, enrichment of PsychENCODE and GWAS-derived risk variants among PeCE-sensitive genes supports the notation that environmentally induced transcriptional perturbations converge on pathways already implicated in human neurodevelopmental disease. While these associations do not establish direct disease causality, they underscore the translational relevance of the molecular programs identified here and suggest that prenatal vaping may influence developmental trajectories along biologically vulnerable genetic axes.

In summary, our multi-modal analyses reveal that prenatal e-cigarette exposure induces coordinated transcriptional, metabolic, synaptic, and developmental reprogramming across the neonatal brain, with particularly strong vulnerability in the developing striatal and cortical regions. We propose that disruption of lipid-dependent developmental homeostasis creates a mismatch between metabolic adaptation and structural maturation, thereby delaying striatal synaptic and cortical development and assembly. PeCE induces broad molecular signatures consistent with elevated Ca^2+^ influx capability and dopaminergic responses, coupled with a shift toward lipid-based energy production in the CPu that may uncouple the energy production from structural maturation. This imbalance manifests as an accumulation of silent synapses and delay in D1MSN developmental progression, while the cortex fails to sustain cholesterol biosynthesis during a critical window of circuit assembly. Together, our findings provide a mechanistically informed framework for understanding how maternal vaping may increase offspring susceptibility to later neurodevelopmental and psychiatric disruptions, which highlight prenatal e-cigarette exposure as an environmental factor warranting closer developmental investigation.

## Materials and methods

### Prenatal e-cigarette exposure animal model

All procedures and protocols for animal experiments were approved by the Institutional Animal Care and Use Committee of Loma Linda University and followed the guidelines in the National Institutes of Health Guide for the Care and Use of Laboratory Animals.

A chronic intermittent e-cigarette (CIEC) exposure rat model was as previously described^20^. Pregnant Sprague-Dawley rats were purchased from Charles River Laboratories (Portage, MI) and were randomly divided into two groups that were exposed to either e-cigarette aerosol (2.4% nicotine, BluPlus Cig) or control air. E-cig aerosol was delivered with a puff duration of 4 second, 3 puffs in an inter-puff interval of 30 seconds per vaping episode, and one episode per 1 hour in the dark phase of 12 hours each day, which generates nicotine blood pharmacokinetics in the pregnant rats similar to those observed in human e-cigarette users^20, 71, 72^. The dams were exposed to e-cigarettes for a total of 17 days from gestational day 4 (E4) to E20.

### Brain tissue harvest and cryosection

Postnatal day 7 (P7) offspring from both prenatal e-cigarette exposure and control rats were used for spatial transcriptomic experiments. P7 rat pups were euthanized under deep isoflurane anesthesia. The brains were harvested and embedded in cold optimal cutting temperature (OCT) medium and snap frozen in isopentane on dry ice. Tissue blocks were equilibrated at -14 °C for at least 30 minutes before coronal cryo-sectioning at a thickness of 10 μm using a CryoStar NX70 (Epredia). For each brain, two sections (one anterior and one posterior) were collected and mounted onto spatially barcoded Visium Gene Expression slides (10x Genomics). To ensure that sections from different brains were obtained from comparable regions, we established a postnatal day 7 (P7) rat brain reference map by performing serial sectioning of the P7 rat brain at 50 μm intervals, followed by H&E staining. This allowed for the assessment of neuroanatomical structure and the selection of appropriate sections. The anterior section was selected based on the following criteria: approximately 0.95 mm anterior to the point where the hippocampus disappears, the size of the anterior commissure and its position relative to the lateral ventricle, and the position of lateral ventricle. The posterior section was chosen based on the shape of the dentate gyrus and the angle formed by the bottom of CA3 and dentate gyrus. Sections were immediately processed following the 10x Genomics Visium Gene Expression protocol (CG000239 Rev F User Guide Visium Spatial Gene Expression Reagent Kits).

### Spatial transcriptomic library preparation and sequencing

34 sections were collected onto the 10x Visium Gene Expression slides and processed according to the manufacturer’s instructions. Briefly, the slides were fixed in cold methanol at -20 °C for 30 minutes and stained with H&E. Brightfield images of the H&E-stained sections were acquired at 10X magnification using an Olympus APX100 Microscope with an auto-stitching function. Tissue permeabilization was performed for 18 minutes, as determined by the tissue optimization kit. After reverse transcription and second strand synthesis, cDNA was amplified through 14 cycles of PCR, followed by fragmentation, end repair and A-tailing, and cleanup and adaptor ligation. A 14-cycle sample index PCR was then performed, followed by double-sided size selection using SPRI beads. Library quantification was conducted using Qubit 4.0 (Life Technologies), and quality control was assessed using a TapeStation 2200 (Agilent Technologies). Sequencing libraries were prepared at the Center for Genomics at Loma Linda University (LLU). The pooled libraries were sequenced on an Illumina NextSeq 2000 (LLU Center for Genomics) and NovaSeq X (UCLA Neuroscience Genomics Core) with paired-end sequencing (Read 1: 28 bp, i7 index: 10 bp, i5 index: 10 bp, Read 2: 90 bp). Detailed sequencing depth and quality information are listed in **Supplementary Data 1**.

In parallel, we conducted snRNA-seq on a subset of control samples derived from 100 μm thickness tissue adjacent to the Visium spatial sections to generate a matched snRNA-seq dataset (**Fig. 1b**). Overall, 23,022 nuclei adjacent to the anterior Visium sections and 61,405 nuclei adjacent to the posterior Visium sections passed quality control filtering and were included for analysis.

### snRNA-seq library preparation and sequencing

1 mm diameter tissue puncture from CPu and NAc regions from P7 male rats were collected and pooled into control and e-cigarette groups, respectively (n = 5 for each group). Single nucleus isolation was performed following the 10x Genomics protocol (CG000124 Isolation of Nuclei for Single Cell RNA Sequencing). Library preparation for snRNA-seq was conducted according to the manufacturer’s instructions. Based on the cell suspension volume calculator table supplied by 10x Genomics, 10,000 nuclei were loaded into the Chromium Controller along with barcode-beads and reverse transcription (RT) reagents to generate single nucleus Gel Bead-in-Emulsions (GEMs). Following cDNA amplification and cleanup, library construction was performed through cDNA fragmentation, end repair and A-tailing, cleanup, adaptor ligation, and sample index PCR. Library quantification was conducted using Qubit 4.0 (Life Technologies), and quality control was assessed using a TapeStation 2200 (Agilent Technologies). Sequencing libraries were prepared at the Center for Genomics at Loma Linda University (LLU). The pooled libraries were sequenced on an Illumina NextSeq 2000 (LLU Center for Genomics) with paired-end sequencing (Read 1: 28 bp, i7 index: 10 bp, i5 index: 10 bp, Read 2: 90 bp).

### RNAScope *in situ* hybridization and imaging

The P7 rat pups were euthanized under deep isoflurane anesthesia. The whole brain was harvested and embedded in cold optimal cutting temperature medium (OCT) and snap frozen in isopentane on dry ice. 10 μm coronal sections close to the anterior or posterior Visium sections were cut, mounted on Superfrost Plus slides and stored at – 80 °C until use. *in situ* hybridization was performed following the manual for the ACD RNAscope Fluorescent Multiplex Assay. Fresh frozen slides were removed from -80 °C storage and immediately transferred to prechilled 4% paraformaldehyde for 1 hour at 4 °C. After dehydration through a graded ethanol series, the sections were incubated in RNAScope Hydrogen Peroxide for 10 min at room temperature, followed by washing steps in distilled water. The sections were incubated in RNAScope Protease IV reagent for 30 min at room temperature. After washing in distilled water, probes were added to hybridize with separate sections in a HybEZ oven for 2 h at 40 °C. The following probes were used: *Cartpt* (Cat# 449511-C1), *Pnoc* (Cat#510521-C2), *Nr4a3* (Cat# 1559351-C1), *Gpr52* (Cat# 314341-C1), RNAScope negative control (Cat# 320871), and RNAScope positive control (Cat# 320891). After probe incubation, the slides were washed twice in 1x wash buffer. The probe-specific signal was developed by sequential hybridization of amplification reagents AMP1-3 at 40 °C for 30-30-15 min, respectively. Slides were incubated with HRP-C1 at 40 °C for 15 minutes and washed twice and re-incubated with TSA vivid fluorophore 520 for labeling the C1 probe at 40 °C for 30 minutes. For sections staining with two different probes, the slides were incubated with HRP blocker at 40 °C for 15 minutes after a 2x wash. Wash and incubation were repeated by using HRP-C2, diluted TSA vivid fluorophore 570 for labeling C2, and HRP blocker with 15-30-15 minutes of incubation, respectively. Slides were counterstained with DAPI (1:10,000 dilutions) for 1 minute followed by washing and mounting with Prolong Gold Antifade Mountant (Invitrogen, P36930). The RNAScope signal was imaged at 10X magnification using an Olympus APX100 Microscope with an auto-stitching function.

### Sample processing and data analysis of lipidomics

For homogenized tissue, 20-100 mg of tissue were collected in a 2 ml homogenizer tube pre-loaded with 2.8mm ceramic beads (Omni #19-628). 0.75mL PBS was added to the tube and homogenized in the Omni Bead Ruptor Elite (3 cycles of 10 seconds at 5 m/s with a 10 second dwell time). Homogenate containing 1-8mg of original tissue was transferred to a glass tube for extraction. A modified Bligh and Dyer extraction^29^ was carried out on all samples. Prior to biphasic extraction, a standard mixture of 74 lipid standards based on Avanti Ultimate Splash ONE mix (Avanti 330820) with Splash Booster (Avanti 330740) was added. Following two successive extractions, pooled organic layers were dried down in a Thermo SpeedVac SPD300DDA using ramp setting 4 at 35 degrees C for 45 minutes with a total run time of 90 minutes. Lipid samples were resuspended in 1:1 methanol/dichloromethane with 10mM Ammonium Acetate and transferred to robovials (Thermo 10800107) for analysis.

Samples were analyzed on the Sciex 6500+ with DMS device with an expanded targeted acquisition list consisting of 1450 lipid species across 17 subclasses. Differential Mobility Device was tuned with EquiSPLASH LIPIDOMIX (Avanti 330731). Data analysis performed on an in-house data analysis platform comparable to the Lipidyzer Workflow Manager^73^. Quantitative values were normalized to mg of tissue.

## Data processing

### Visium spatial transcriptome analysis

FASTQ files and histology images were processed with the Space Ranger software v.2.0.1 and mapped against the Cell Ranger rn7 reference genome cellranger6_GEX_rn7 to generate raw count data. QC summaries after pipeline processing are available in Supplementary Data 1. Seurat (v.5.0.1) and R (v.4.3.2) were used for downstream data analysis. Visium spots with fewer than 500 genes were excluded. Spots with nCount_Spatial < 200 as well as more than 15% mitochondrial genes were filtered out. We followed the standard Seurat integration workflow^74^ by identifying pairwise anchors (top 3000 most variable features) across samples. For each group (male anterior, female anterior, male posterior, female posterior), 8 or 9 Visium datasets were integrated by IntegrateData() with normalization.method = “SCT”, generating one combined Seurat object for each group followed by the standard Seurat clustering workflow. Clustering and visualization were performed using Uniform Manifold Approximation and Projection (UMAP) as the dimensionality reduction algorithm. To identify region-specific transcriptomic changes induced by prenatal e-cig exposure, we used the MAST package^28^ (v.1.28.0) to perform differential gene expression analysis with the following criteria: test.use = “MAST”, min.pct = 0.2, logfc.threshold = 0.25, p_val_adj < 0.05.

### Generation of single-nucleus transcriptomic count matrix

Raw sequencing data were processed with Cell Ranger (v7.2.0) for sample demultiplexing, cell barcode detection, and single-nucleus expression count matrix generation. The ‘cellranger mkfastq’ command was used to demultiplex the samples, extract the UMI barcodes, and generate the FASTQ files. The gene expression count matrices for each sample were generated using the ‘cellranger count’ command by aligning the reads to the rn7 genome.

### Single-nucleus transcriptomic data analysis

Seurat (v.5.0.1) and R (v.4.3.2) were used for data analysis. Cells with fewer than 300 genes were excluded. Cells with many uniquely expressed genes (≥98% percentile) as well as more than 15% mitochondrial genes were filtered out. After filtering, 22,278 nuclei from CPu and 14,414 nuclei from NAc were merged into one Seurat object. The gene expression profile of each single nucleus was normalized by log normalization - NormalizeData(), followed by scaling data - ScaleData() - to adjust for highly variable genes. The top 2000 most highly variable genes were selected using the Seurat function ‘FindVariableGenes’ with ‘vst’ method. The variable genes were used to calculate the top 30 principal components, and the cells were then classified into clusters with the Seurat ‘FindClusters’ function.

### Mapping and quantification of bulk RNA-seq data from CPu and mPFC tissues

Raw paired-end RNA-sequencing reads (151 bp) generated from caudoputamen (CPu) and medial prefrontal cortex (mPFC) tissues were aligned to the rat reference genome (Rn7 assembly) using STAR (v2.7.10b) (PMID: 23104886). Prior to alignment, a STAR genome index was generated from the Rn7 reference genome. Reads were mapped with STAR using the following command: STAR --runThreadN --genomeDir --readFilesIn --readFilesCommand --outFileNamePrefix --outSAMtype SortedByCoordinate - -quantMode.

Aligned BAM files were subsequently used for gene-level expression quantification with StringTie (v2.2.1) (PMID: 25690850). Expression estimation was performed in reference-guided mode using the Ensembl gene annotation for rat (Rattus_norvegicus.mRatBN7.2.113.chr.gtf). StringTie was run on assembly to annotated transcripts only and generating both gene-level abundance tables and Ballgown-compatible output files. The StringTie command “stringtie input.bam -p -G Rattus_norvegicus.mRatBN7.2.113.chr.gtf -e -B -A gene_abundance.txt -o output.gtf. Resulting gene-level expression estimates were used for downstream differential expression and comparative analyses.

### Differential gene expression and gene set enrichment analysis

Gene-level expression estimates obtained from StringTie were input to DESeq2^75^ to perform differential gene expression analysis between control and e-cigarette within CPu and mPFC tissues. Genes were tested for differential expressions between groups, and resulting statistics included log2 fold changes and associated significance values. For pathway-level interpretation, genes were ranked based on the signed differential expression statistic derived from the DGE analysis (log2 fold change weighted by statistical significance). This ranked gene list was used as input for pre-ranked Gene Set Enrichment Analysis (GSEA)^76^. Enrichment analyses were performed for Gene Ontology (GO) biological process terms and for curated Hallmark gene sets^77^, enabling identification of coordinated transcriptional programs associated with each condition. Enrichment scores and normalized enrichment scores were computed using phenotype-independent permutations, and statistical significance was assessed with multiple-testing correction. Enriched pathways were interpreted based on directionality and consistency with observed differential expression patterns.

### Module score calculation for mechanistic gene groups

To assess coordinated activity of predefined mechanistic gene programs, curated functional gene groups representing key components of calcium influx (*Cacna1c, Cacna1d, Cacna1e, Cacna2d1, Grin1, Grin2a, Grin2b*) and dopamine signaling regulators (*Drd1, Drd2, Adcy1, Ppp1r1b, Gnal, Rgs9, Adcy5, Prkaca, Prkacb, Creb1, Atf2, Ppp1ca, Ppp2ca*) were defined a priori based on known biology^78, 79, 80^. Module scores for each gene group were computed at spot level across each spatial region using the AddModuleScore function implemented in Seurat^81^. Module scores were calculated independently for each functional gene group at the spot level and computed the mean and median scores across each spatial region in control and e-cigarette samples. Seurat calculates the average normalized expression of genes in each spot and subtracts the aggregated expression of control gene sets matched for overall expression level, thereby controlling technical effects and gene expression magnitude.

### Mapping the prenatal e-cig exposure induced DEGs with the human neurodevelopmental disorder data

The PsychENCODE database is a publicly accessible data resource developed by the PsychENCODE Consortium^82^. It provides large-scale multi-omics datasets derived from human brain samples to facilitate the investigation of genetic and molecular mechanisms underlying psychiatric disorders. The database includes transcriptomic data (bulk RNA-seq, snRNA-seq, gene expression across brain regions, development stages, and disease conditions), epigenomic data (ATAC-seq, ChIP-seq, and DNA methylation), genomic data (including GWAS variants, eQTLs, and Hi-C), and associated clinical metadata. We downloaded differentially expressed genes (DEGs) and GWAS SNPs from human brain PsychENCODE data^22, 23^. All human data were converted to the rat (rn7) genome using a lift-over approach for comparative analysis. The DEGs were derived from snRNA-seq datasets related to three neurological disorders: autism spectrum disorder (ASD), bipolar disorder, and Schizophrenia. GWAS SNPs were uplifted to the rat rn7 genome. We intersected the orthologous SNP locations with DEGs and non-DEGs and constructed a two-by-two contingency table for Fisher’s exact test which was used to determine whether orthologous SNPs were enriched in orthologous DEGs compared to non-DEGs.

Human gene sets were downloaded from Coe et al. ^34^ (COE), Kaplanis et al. (DDD)^35^ and SFARI gene (https://gene.sfari.org/database/human-gene/) and then mapped to rat orthologs using biomart database. Enrichment testing was performed using fisher.test() and P-values were Bonferroni corrected for multiple tests using p.adjust() in R. Odds ratio heatmap was plotted using ggplot2.

## Acknowledgments

This work was funded in part by the National Institutes of Drug Abuse (NIDA) grant 1U01DA058278 (CW), National Institutes of Health (NIH) grant S10OD019960 (CW), Ardmore Institute of Health grant 2150141 (CW), and Dr. Charles A. Sims’ gift to LLU Center for Genomics. Partial schematic illustrations in Figure 1 were generated using Biorender. We appreciate Dr. Pingyu Zeng’s technical support for RNAScope.

## Author contributions

Conceptualization and supervision: CW; sample collection: WC, ZC and YL; animal exposure: Y L and DX; data generation: WC and ZC; methodology: WC, ZC, CN; data analysis and visualization: WC, ZC, CN and XP; writing – original draft: WC, ZC, CN; writing – review and revision: CW, WC, ZC, YD, MM, WJ, WXS. Manuscript final revision: CW.

---

**Supplementary Data 1.** Visium spatial transcriptomics library sequence quality data summary.

**Supplementary Data 2.** Region-specific genes across anterior brain region in control P7 rat (Fig. 1g)

**Supplementary Data 3.** Region-specific genes across posterior brain region in control P7 rat (Fig. 1h)

**Supplementary Data 4.** Differential expressed genes from anterior and posterior Visium spatial transcriptomics dataset

**Supplementary Data 5.** Overlapping DEGs between male and female within each subregion from anterior Visium spatial transcriptomics dataset

**Supplementary Data 6.** Overlapping DEGs between male and female within each subregion from posterior Visium spatial transcriptomics dataset

**Supplementary Data 7.** Fig. 2b Region-specific DEG list for P7 anterior rat brain

**Supplementary Data 8.** Fig. 2c Region-specific DEG list for P7 posterior rat brain

**Supplementary Data 9.** E-cig induced DEGs from P7 Visium data overlapping with ASD DEGs

**Supplemental Figure 1.**
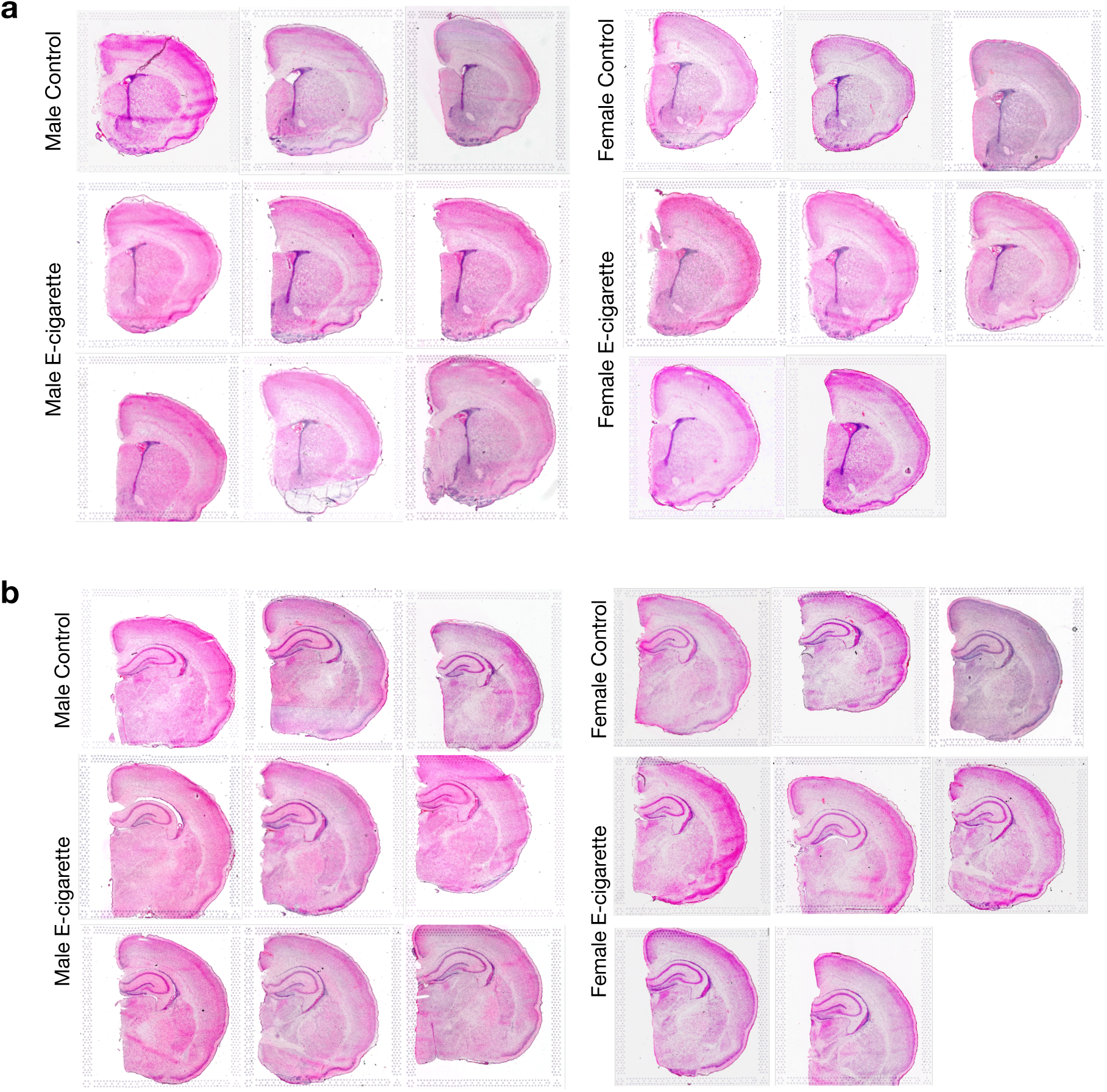
H&E staining of all the Visium anterior and posterior sections. H&E staining of all the 34 Visium spatial transcriptomic sections from postnatal day 7 rat brains; anterior (a) and posterior (b) sections.

**Supplemental Figure 2.**
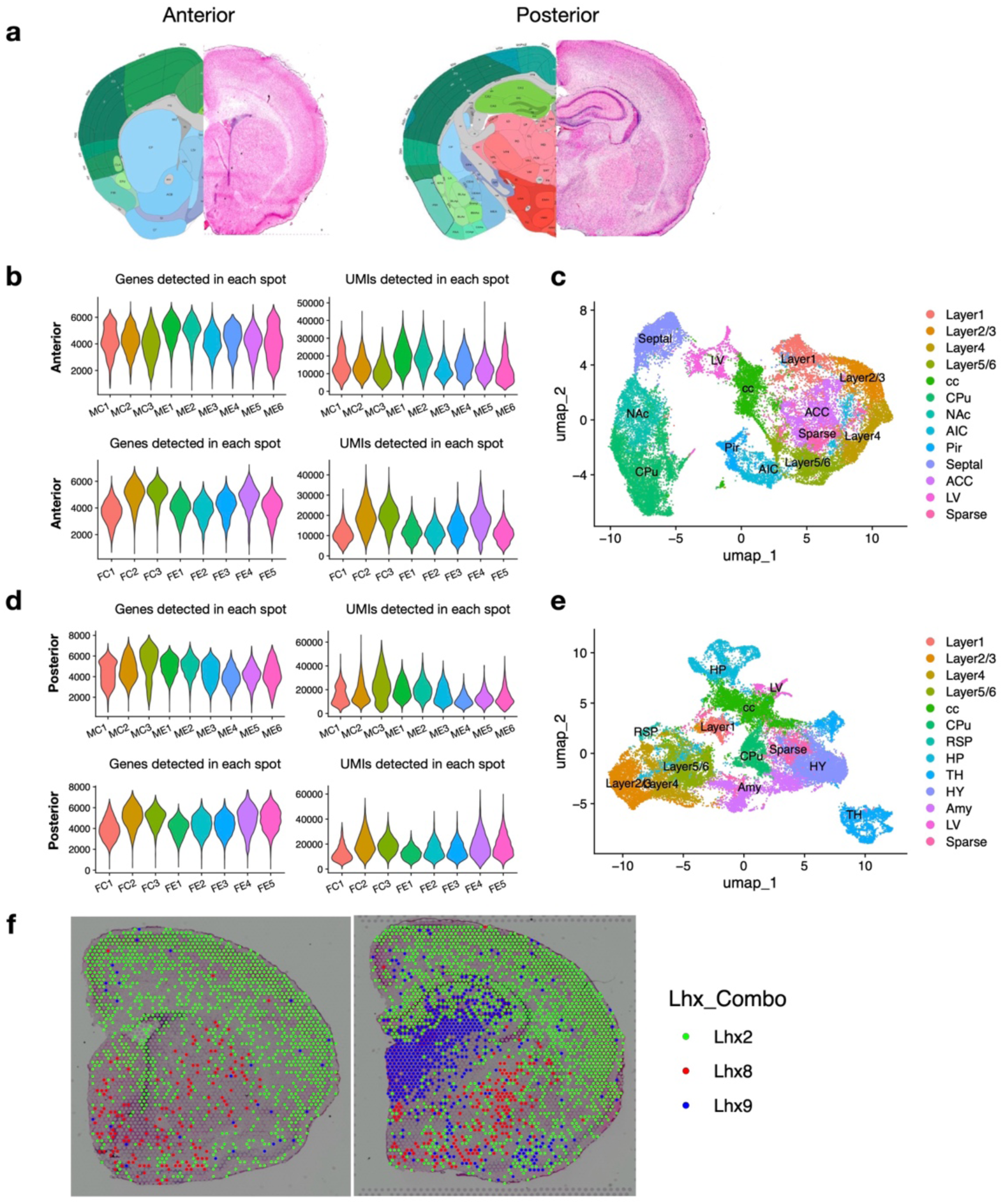
Quality control of Visium spatial transcriptomic data. **a.** The brain pictures on the left side of the images were taken from Allen Mouse Brain Atlas as brain region annotation references. On the right side are representative H&E stained anterior and posterior sections used for generating Visium spatial transcriptomics. **b.** Violin plots showing the number of genes expressed and UMI counts per Visium spot in all the anterior spatial transcriptomic datasets**. c.** UMAP visualization of 28,055 Visium spots from male postnatal day 7 anterior regions. **d.** Violin plots showing the number of expressed genes and UMI counts per Visium spot in all the posterior spatial transcriptomic datasets. **e.** UMAP visualization of 31,541 Visium spots from male postnatal day 7 posterior regions. **f.** Spatial expression of Lhx2, Lhx8, and Lhx9 on the P7 male anterior (left) and posterior (right) brain.

**Supplemental Figure 3.**
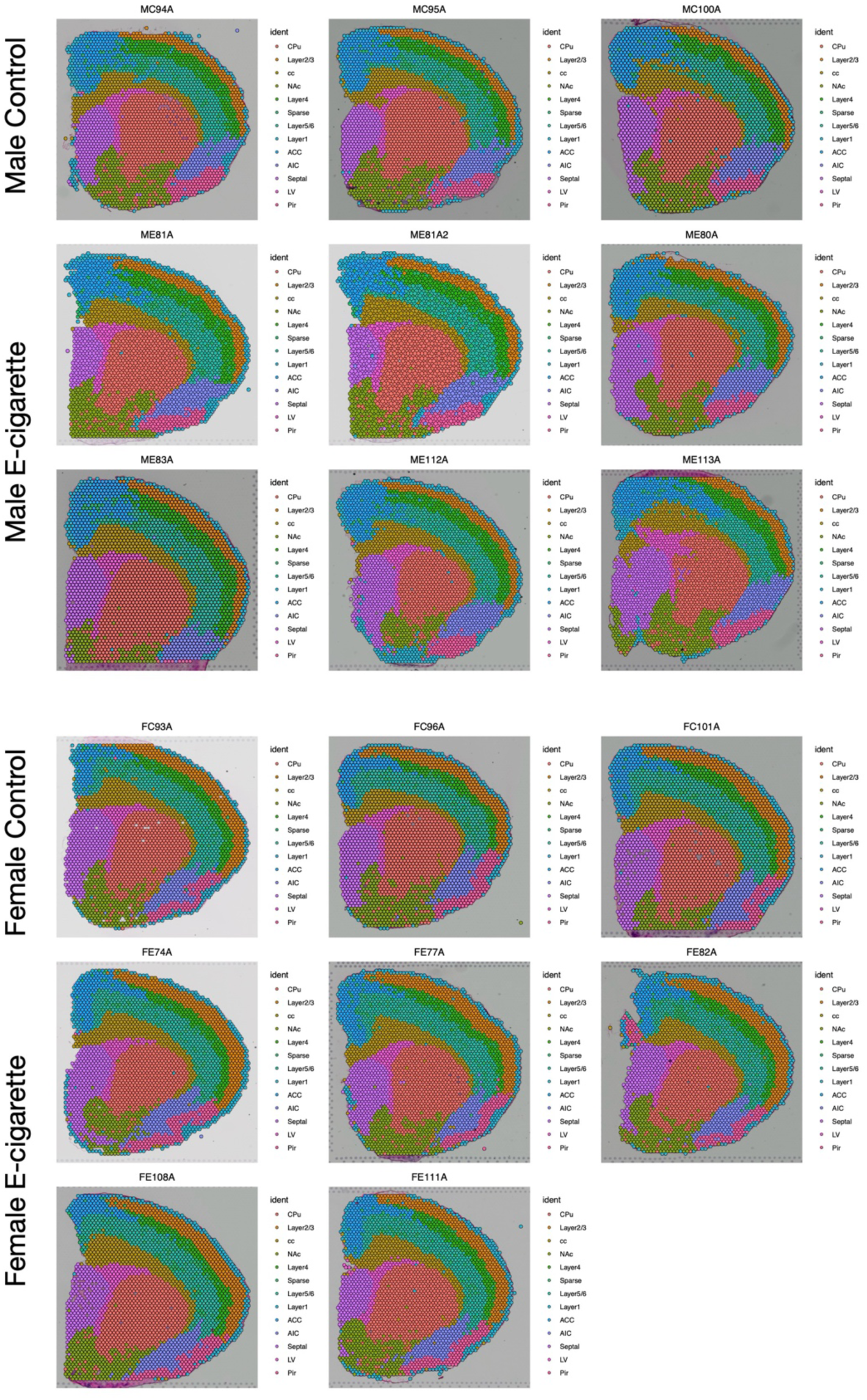
Graph-based unsupervised cluster identification of all 17 anterior sections. Each spot is colored based on the transcriptional signature using the Louvain clustering algorithm.

**Supplemental Figure 4.**
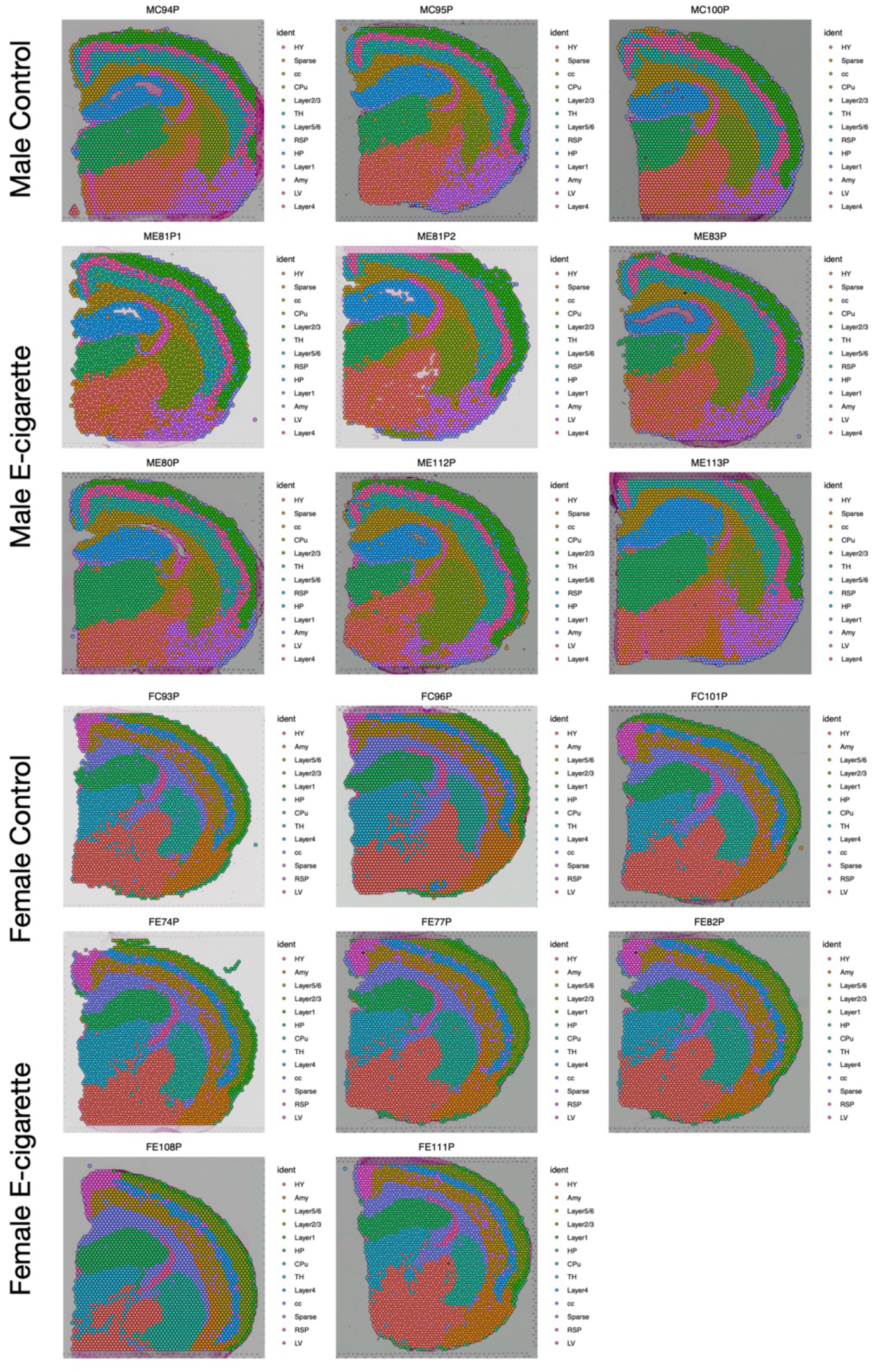
Graph-based unsupervised cluster identification of all 17 posterior sections. Each spot is colored based on the transcriptional signature using the Louvain clustering algorithm.

**Supplemental Figure 5.**
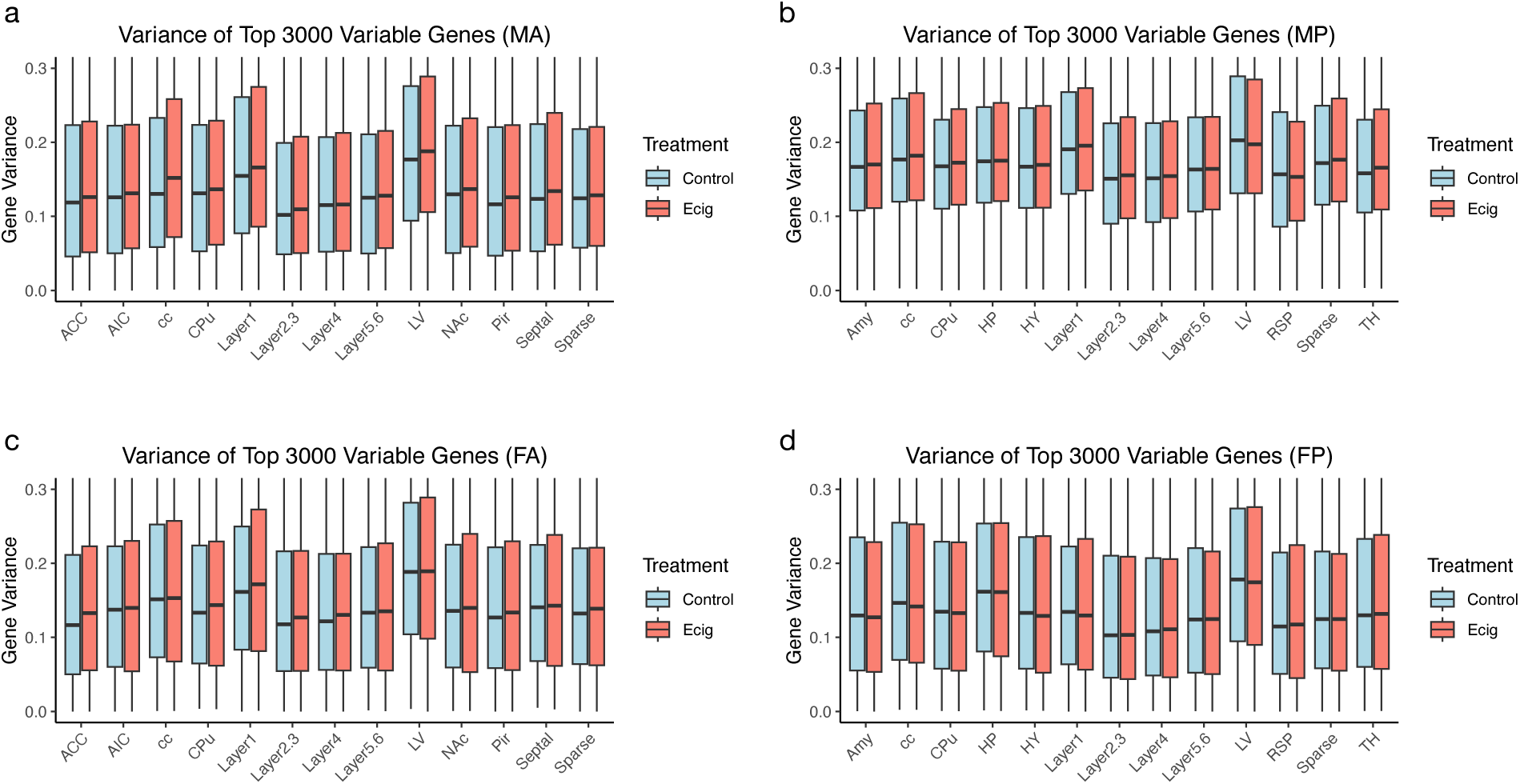
The variance of the top 3000 most variable genes in Visium spatial transcriptomic datasets. **a.** Boxplot showing the variance of the 3000 most variable genes in the male anterior (MA) Visium dataset in each subregion. **b.** Boxplot showing the variance of top 3000 most variable genes in the male posterior (MP) Visium dataset in each subregion. **c.** The boxplot showing the variance of the 3000 most variable genes in the female anterior (FA) Visium dataset in each subregion. **d.** Boxplot showing the variance of the 3000 most variable genes in the female posterior (FP) Visium dataset in each subregion. MA, male anterior; FA, female anterior; MP, male posterior, FP, female posterior.

**Supplemental Figure 6.**
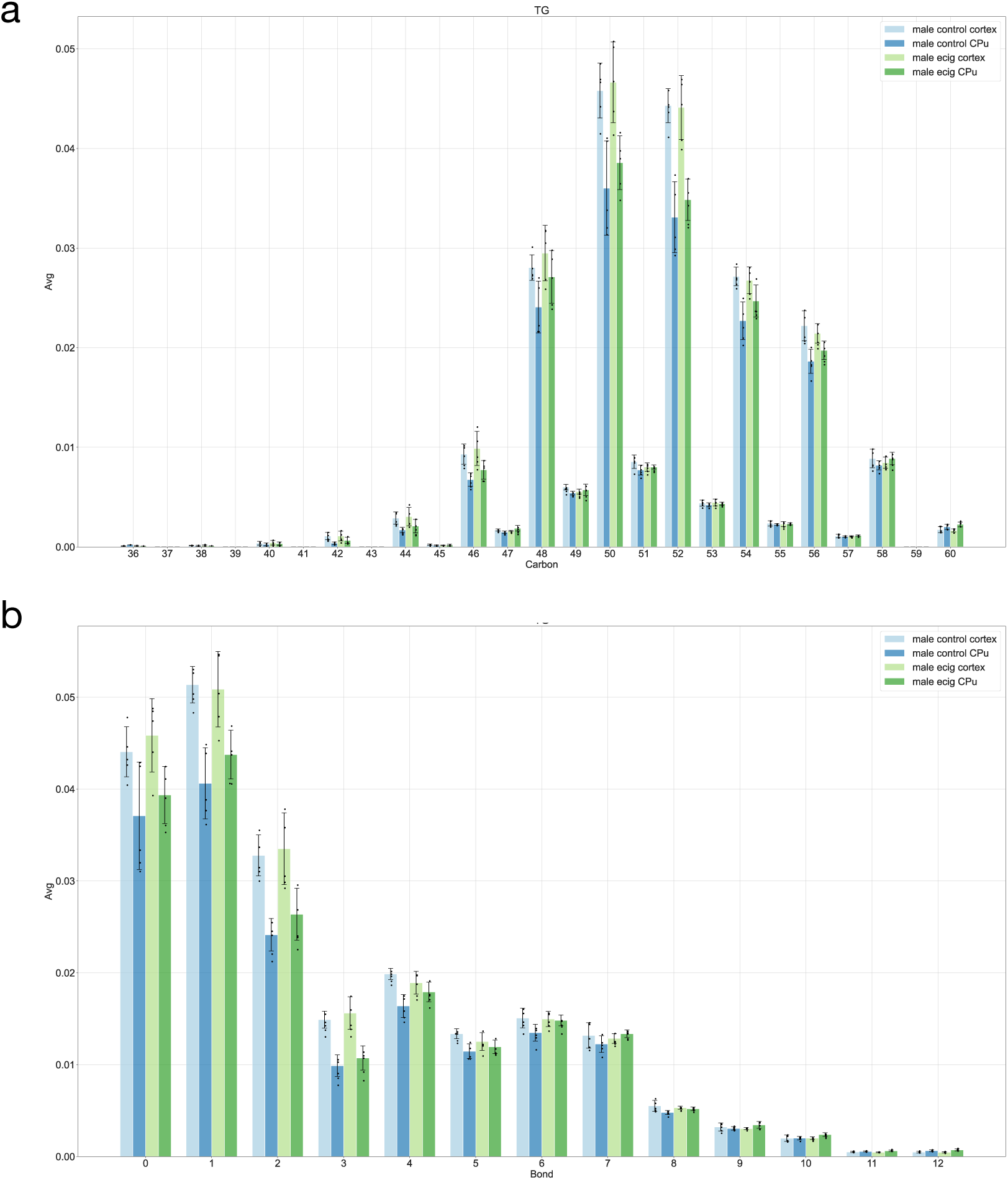
Lipidomics analysis showing the level of triacylglycerols in the cortex and CPu regions from control and e-cigarette-exposed P7 rats. The data were visualized by carbon chain length (a) and saturation level (b).

**Supplemental Figure 7.**
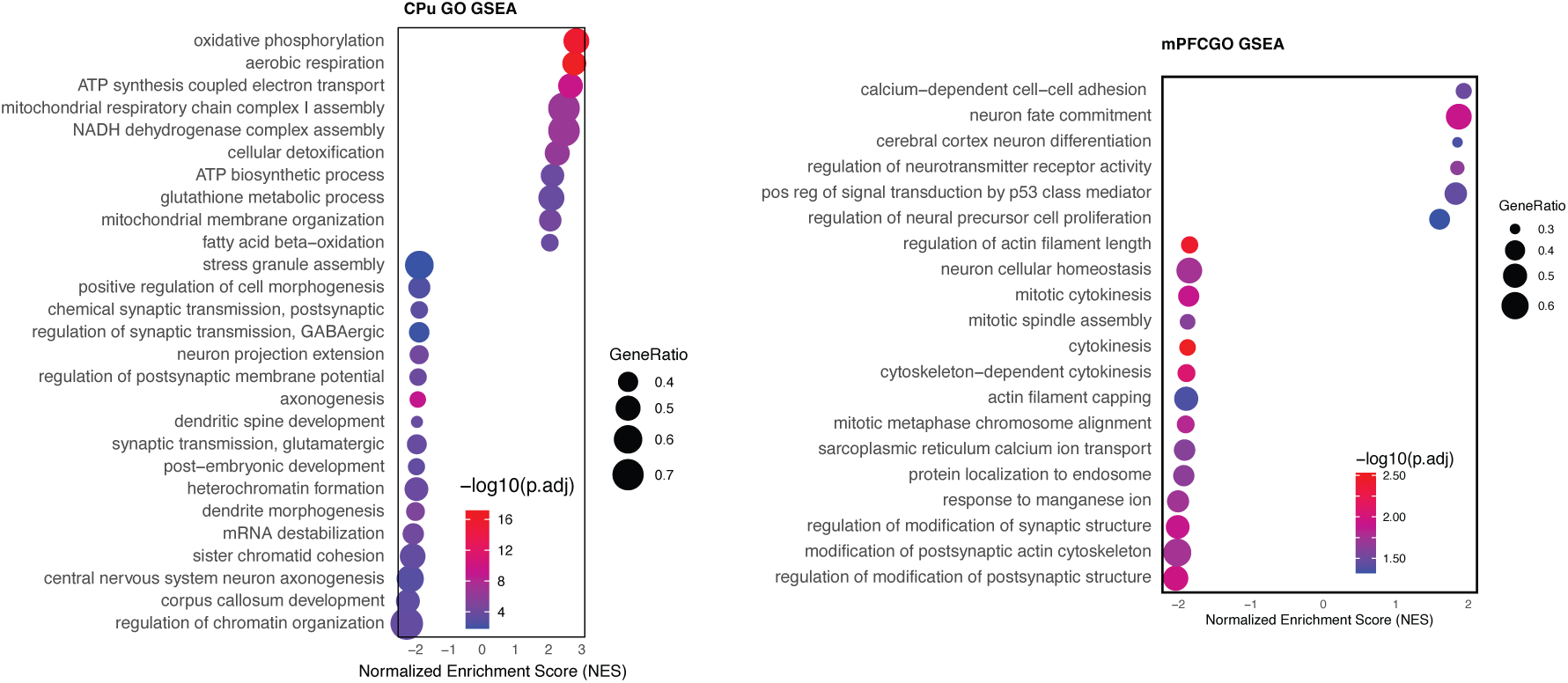
Dot plot showing the top Gene Ontology biological process terms enriched based on Gene set enrichment analysis (GSEA) of bulk RNA-seq data in CPu and mPFC.

**Supplemental Figure 8.**
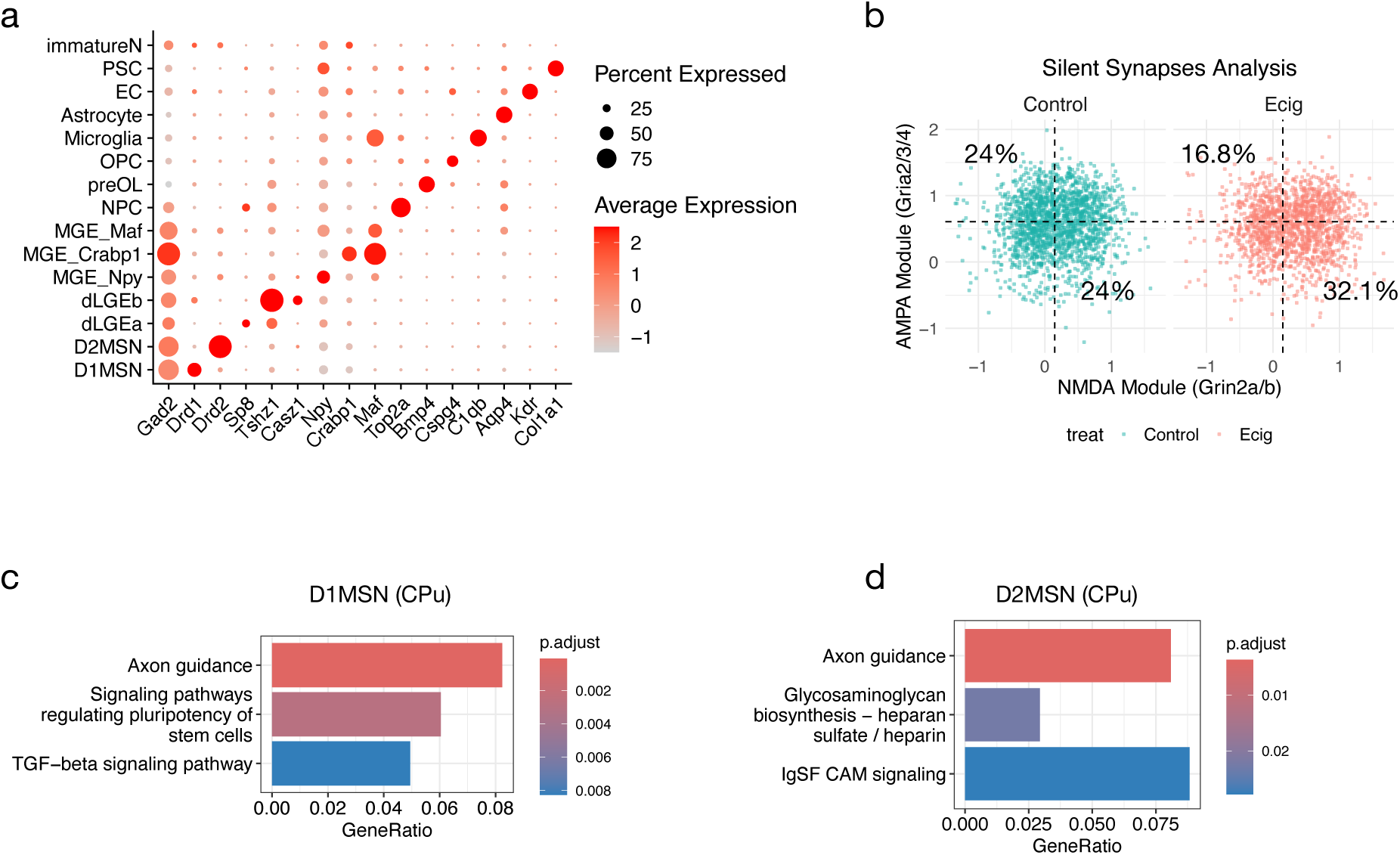
snRNA-seq analyses of P7 CPu and NAc regions. **a.** Dot plot showing representative marker gene expression across 15 cell clusters derived from CPu and NAc of P7 rat brain. **b.** Scatter plot showing the module score of AMPA and NMDA of CPu D2MSN in four quadrants. **c-d.** Top enriched KEGG pathways in D1MSN (c) and D2MSN (d) from CPu, respectively. Down-regulated DEGs were used for the analysis.

